# Infiltrating lipid-rich macrophage subpopulations identified as a regulator of increasing prostate size in human benign prostatic hyperplasia

**DOI:** 10.1101/2024.06.07.597992

**Authors:** Nadia A. Lanman, Era Meco, Philip Fitchev, Andree K. Kolliegbo, Meaghan M. Broman, Yana Filipovich, Harish Kothandaraman, Gregory M. Cresswell, Pooja Talaty, Malgorzata Antoniak, Svetlana Brumer, Alexander P. Glaser, Andrew M. Higgins, Brian T. Helfand, Omar E. Franco, Susan E. Crawford, Timothy L. Ratliff, Simon W. Hayward, Renee E. Vickman

**Author notes:** **Corresponding Author:** To whom correspondence should be addressed: Renee E. Vickman, PhD, Department of Surgery, Endeavor Health, 1001 University Place, Room 272, Evanston, IL 60201-3137.

## Abstract

Macrophages exhibit marked phenotypic heterogeneity within and across disease states, with lipid metabolic reprogramming contributing to macrophage activation and heterogeneity. Chronic inflammation has been observed in human benign prostatic hyperplasia (BPH) tissues, however macrophage activation states and their contributions to this hyperplastic disease have not been defined. We postulated that a shift in macrophage phenotypes with increasing prostate size could involve metabolic alterations resulting in prostatic epithelial or stromal hyperplasia. Single-cell RNA-seq of CD45^+^ transition zone leukocytes from 10 large (>90 grams) and 10 small (<40 grams) human prostates was conducted. Macrophage subpopulations were defined using marker genes. BPH macrophages do not distinctly categorize into M1 and M2 phenotypes. Instead, macrophages with neither polarization signature preferentially accumulate in large versus small prostates. Specifically, macrophage subpopulations with altered lipid metabolism pathways, demarcated by *TREM2* and *MARCO* expression, significantly accumulate with increased prostate volume. *TREM2*^+^ and *MARCO*^+^ macrophage abundance positively correlates with patient body mass index and urinary symptom scores. TREM2^+^ macrophages have significantly higher neutral lipid than TREM2^−^ macrophages from BPH tissues. Lipid-rich macrophages were observed to localize within the stroma in BPH tissues. *In vitro* studies indicate that lipid-loaded macrophages increase prostate epithelial and stromal cell proliferation compared to control macrophages. These data define two new BPH immune subpopulations, TREM2^+^ and MARCO^+^ macrophages, and suggest that lipid-rich macrophages may exacerbate lower urinary tract symptoms in patients with large prostates. Further investigation is needed to evaluate the therapeutic benefit of targeting these cells in BPH.

## Introduction

Benign prostatic hyperplasia (BPH) is a common age-related disease in men and is a cause of lower urinary tract symptoms (LUTS). LUTS include a range of storage, voiding, and post-micturition issues, evaluated most commonly by the self-reported International Prostatic Symptom Score (IPSS) questionnaire^1^. Increasing prostate size does not directly correlate with IPSS due to the multifactorial causes of LUTS, the variability of overall prostate shape (e.g. presence or absence of a median lobe), varying degrees of bother reported by patients, as well as periurethral fibrosis in some patients^2–4^. As a result, most studies focused on histologic prostate hyperplasia need to include a measurement of prostate weight or volume rather than rely solely on symptom score.

Severe prostatic inflammation has been shown to decrease the therapeutic efficacy of 5α-reductase inhibitors or α-adrenergic blockers in BPH patients^5^. Our recent work indicates that CD45^+^ inflammatory cells accumulate within prostates of increased size^6^, but whether specific immune subpopulations are over-represented in these tissues is unknown. Macrophages and T cells are the dominant leukocytes accumulating in BPH tissues^6,7^. We have also reported that targeting systemic inflammation with tumor necrosis factor (TNF)-antagonists decreases BPH incidence in autoimmune disease patients and decreases macrophage accumulation in prostate tissues^6^. A greater understanding of the inflammatory cell states present in BPH tissues is necessary to elucidate immune-targeted therapies with long-term efficacy in this aged patient population.

Macrophage diversity and plasticity have been observed in numerous diseases^8,9^. Macrophage polarization, or the transition to specific phenotypic states, has previously been categorized into two broad phenotypes: pro-inflammatory M1 or anti-inflammatory M2^10,11^. Since M1 and M2 macrophages are the extremes of a functional cell state continuum, it is not surprising that these classifications do not reflect the broad range of macrophage phenotypic diversity observed *in vivo*. Recently, a variety of macrophage polarization states have been described, including, but not limited to, interferon-inducible cell (IFNIC)^12^, metallothionein (Mac-MT)^13^, and lipid metabolism-dysregulated macrophages identified with marker genes such as triggering receptor expressed on myeloid cells 2 (TREM2) and macrophage receptor with collagenous structure (MARCO)^14,15^.

Lipids impact macrophage function through cytokine release and intracellular metabolism in various conditions^16^. Lipid-loading can lead to activation of peroxisome proliferator activated receptors (PPARs), with downstream suppression of macrophage inflammatory signaling or induction of alternative macrophage activation in some diseases^17,18^. TREM2 is a multifunctional, lipid-binding transmembrane receptor expressed on myeloid cells that can activate tissue repair pathways or exacerbate inflammation, indicating the importance of disease context^19–21^. TREM2^+^ macrophages have been described in both benign and malignant diseases, including metabolic diseases associated with chronic inflammation, however, TREM2^+^ macrophages tend to share a gene signature that contains lipid and cholesterol metabolism genes across these diverse disease states^19^. Signaling induced by TREM2 can also result in the expression of phagocytic receptors and senescence genes in myeloid cells^19,22^. MARCO is a class A scavenger receptor that promotes phagocytic function in macrophages via cytoskeletal formation of pseudopodia^23^. MARCO^+^ macrophages have been associated with lipid uptake and have been shown to improve metabolic disease by sustaining PI3K activity, but also contribute to cancer progression^14,24,25^.

Obesity, or elevated body mass index (BMI), is associated with systemic inflammation and excessive fat depots. Obesity is also linked to an increased risk of LUTS/BPH, which is perhaps due to both hormonal and inflammatory mechanisms^26^. However, the specific molecular mechanisms that link these diseases are not defined. Elevated circulating lipid levels provide an environment where monocytes or macrophages can uptake lipid via scavenger or lipid-sensing receptors, such as TREM2^19,22^. Examining macrophage phenotypes in BPH tissues could reveal cell states that provide a more mechanistic understanding of how environmental context promotes this disease.

The studies herein utilized scRNA-seq to define infiltrating leukocytes in the progressive inflammatory process that has been observed in larger prostate volumes. Macrophage subclustering analysis identified, for the first time, stromal lipid-loaded TREM2^+^ and MARCO^+^ macrophages in BPH tissues, and showed accumulation of these cells in large versus small prostates. These studies correlated these macrophage subpopulations with clinical parameters and evaluated the effect of lipid-loaded macrophages on epithelial and stromal proliferation in vitro. Together, our data suggest a role for TREM2^+^ and MARCO^+^ macrophages in BPH progression and identify a novel inflammatory target for BPH treatment.

## Results

### Myeloid cells increase in abundance as human prostate size increases

In these studies, existing scRNA-seq human prostate transition zone (TZ) leukocyte data^6^ were expanded to provide a balanced set, where CD45+EpCAM-CD200-leukocytes from 10 large (>90 grams) prostate tissues were compared to leukocytes from 10 small (<40 grams) prostate tissues (Figure 1a). Patient characteristics are listed in Supplementary Data S1. Patients with large prostates had significantly higher IPSS values (p=0.0008) compared to patients with smaller prostates, but there was no significant difference in age or body mass index (BMI) between groups (Supplementary Figure S1a-d). Analysis of a larger BPH patient cohort also indicates that patients with large prostates have significantly higher IPSS (p=0.021), but no difference in age or BMI at the time of surgery (Supplementary Figure S1e-g). Visualization of all CD45^+^ cells by scRNA-seq identified a broad spectrum of inflammatory cell types, with T and myeloid cells as the most abundant major classes (Figure 1b). General cell types were determined using highly expressed cluster markers and CITE-seq^27^ analysis (Supplementary Figure S1h-k). Comparing cells from large versus small prostates identified shifts in the relative abundance of cell clusters, with an increase in some myeloid clusters and a decrease in some T cell clusters (Figure 1c-d). These shifts in cell abundance did not appear to be related to skewed contributions from individual patient samples (Supplementary Figure S2a-b). Evaluation of CD11b^+^ myeloid cells, CD8^+^ and CD4^+^ T cells, and CD19^+^ B cells by flow cytometry in these patient samples indicate that increased myeloid cells (p=0.0096) in large versus small samples are the primary contributor to accumulated leukocytes in BPH (Figure 1e, Supplementary Figure S2c-e). Together, these data suggest that myeloid cells are involved in progressive inflammation that is associated with prostatic hyperplasia.

**Figure 1.**
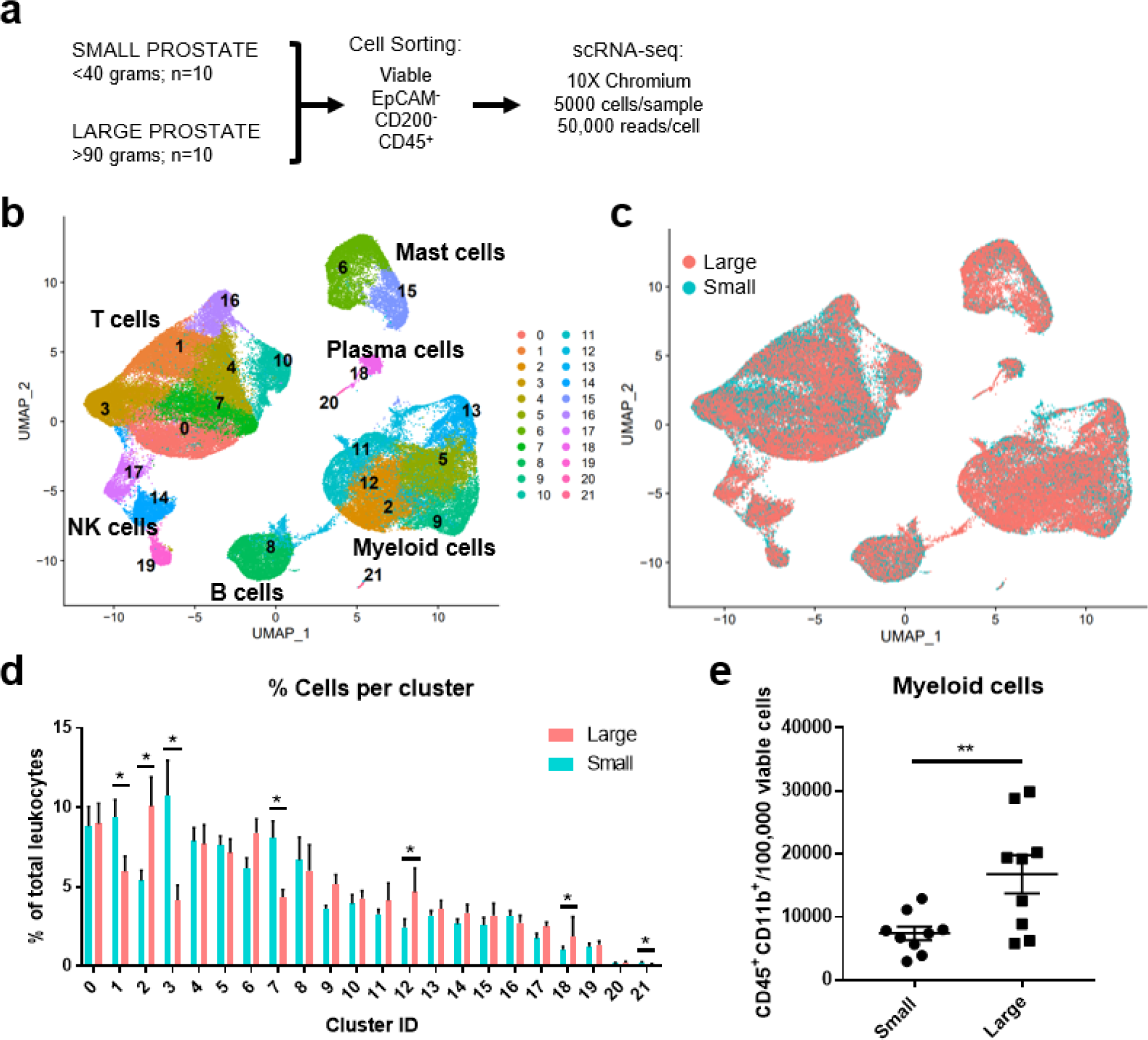
Myeloid cells increase in abundance as human prostate size increases. A total of 10 small (<40 grams) and 10 large (>90 grams) prostate transition zone tissues were digested and prepared for FACS, followed by scRNA-seq of CD45+ cells. **a)** Schematic representing the setup for scRNA-seq studies of human BPH leukocytes. Viable CD45^+^EpCAM^−^CD200^−^ cells were sorted by FACS. Single-cell RNA-seq was conducted using the 10X Chromium system, aiming for 5000 cells/sample at a depth of 50,000 reads/cell. **b)** Uniform manifold approximation and projection (UMAP) plot of 100,459 individual cells from 20 patient samples, demonstrating dominant T and myeloid cell populations. Each color indicates a unique cell cluster. **c)** UMAP of CD45+ cells from the 20 TZ tissues in (a), colored to highlight cells from small (blue) or large (pink) prostates. **d)** Graph representing the mean +/− SEM of the percentage of cells in each scRNA-seq cluster among total CD45^+^ cells between samples in large (pink) versus small (blue) prostates (n=10/group). Asterisks indicate significantly increased (clusters 2, 12, 18) or decreased (clusters 1, 3, 7, 21) populations in large vs small prostate tissues by permutation test (*=FDR <0.05 for each). **e)** Flow cytometry analysis of the number of CD11b+ myeloid cells per 100,000 viable cells in the digested patient samples used for scRNA-seq analysis (available for n=9 per group). **p=0.0096 using an unpaired T test.

### TREM2^+^ and MARCO^+^ macrophage subtypes accumulate in large versus small prostates

Subclustering analysis of prostatic myeloid cells from all 20 patients was performed to investigate specific changes in the subpopulations. Myeloid subclustering analysis demonstrates 14 bioinformatically distinct subclusters (Figure 2a). SingleR does not identify specific subpopulations (Supplementary Figure S3a), so empirical evaluation of biomarkers was used to classify subclusters (Figure 2a-b, Supplementary Figure S3b). Clusters 8, 9 (*FCN1*^+^*VCAN*^+^*S100A8/9*^+^), and 12 (*FTH1*^+^*S100A4*^+^) indicate a monocyte/macrophage phenotype^13^. Clusters 6 and 10 (*CD1C*^+^*FCER1A*^+^*CLEC10A*^+^) were identified as type 2 conventional dendritic cells (cDC2s). Mitochondrial genes are among the top markers for cluster 3 despite normalization. Apparent resident macrophages (Mϕs) were also discovered, with clusters 4 and 5 expressing *LYVE1*, *F13A1*, and *FOLR2*^28^, while clusters 0 and 5 express *C1QC*, *CD74*, and *APOE*^29^. To further substantiate resident populations, alignment of yolk-sac derived macrophage markers using Garnett^30^ indicates this phenotype primarily coincides with clusters 3 and 4, and, to a lesser extent, cluster 5 (Supplementary Figure S4a-b). Cluster 0 is the only cluster with high expression of *TREM2*, previously defined as lipid-associated macrophages in other diseases, such as obesity^31^ and atherosclerosis^32,33^. Cluster 7 contains *TNF*^+^*IL1B*^+^ inflammatory macrophages, but clusters 1 and 2 may also have inflammation-regulating properties since they have upregulated expression of *NR4A1*^+^*SPP1*^+^*TNF*^+^ and *SPP1*^+^*IL1B*^+^ cluster markers, respectively. Minor populations include metallothionein-expressing macrophages (MAC-MT^13^) and interferon-inducible cell macrophages (IFNIC Mϕs^34,35^) in clusters 11 (*MT1F*^+^*MT1E*^+^*MT1X*^+^) and 13 (*ISG15*^+^*MX1*^+^*IFITM3*^+^*IFITM1*^+^), respectively. Permutation analysis of macrophage subclusters indicates a significant decrease in the abundance of clusters 2 (Mϕs), 6 (cDC2), 9, and 12 (monocytes/macrophages), but a significant increase in the abundance of clusters 0 (*TREM2*^+^) and 5 (*MARCO*^+^) in large versus small prostate TZ tissues (Figure 2c-d). These data suggest either proliferation or increased recruitment/polarization of this macrophage phenotype in BPH patients. Evaluation of proliferating cells by aligning myeloid cells to a cell cycle-related gene classifier indicates that, indeed, *TREM2*^+^ and *MARCO*^+^ macrophages (clusters 0 and 5, respectively) are the two myeloid subclusters with highest expression of cell cycle-related genes in BPH (Supplementary Figure S4c-d).

**Figure 2.**
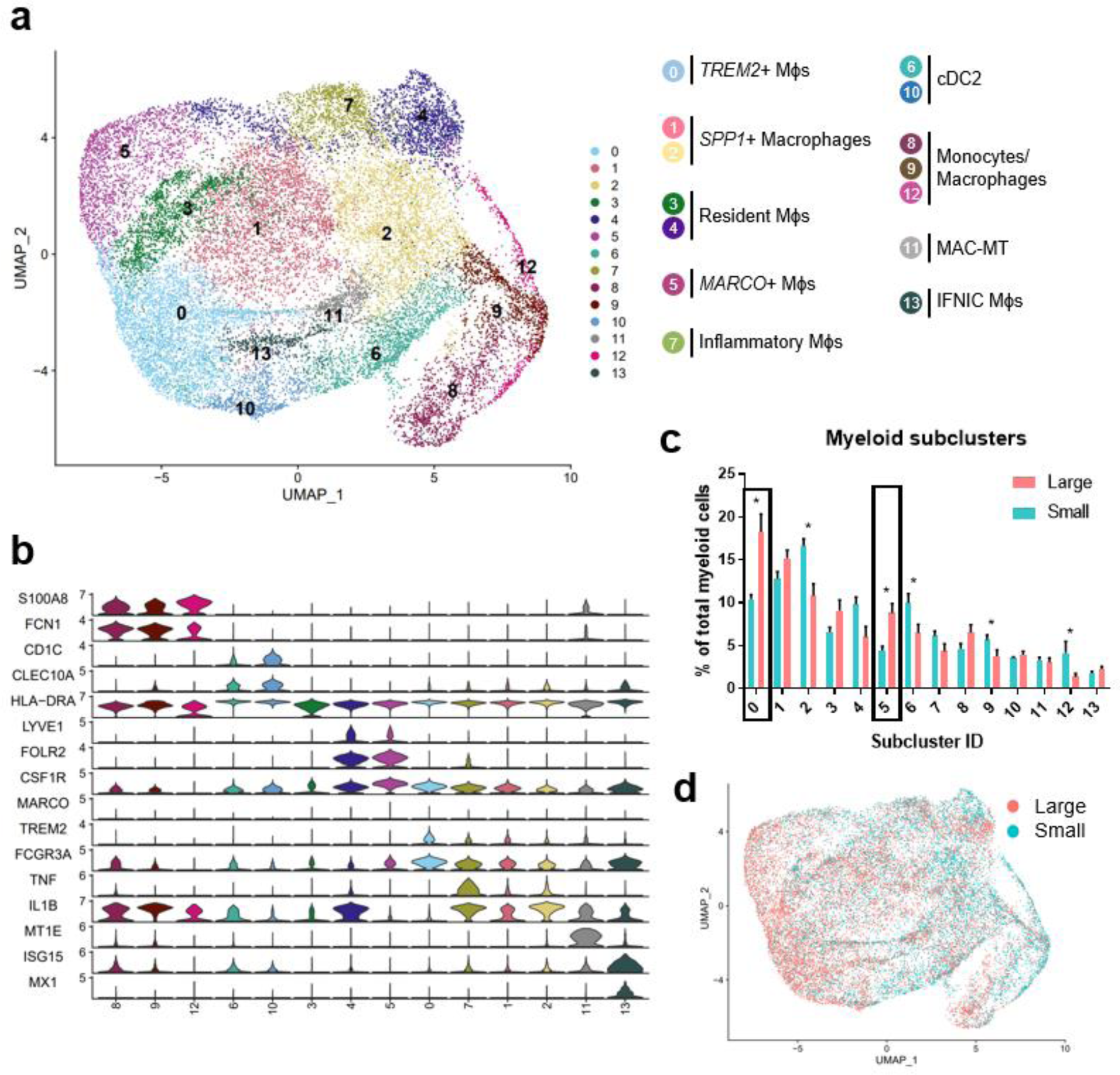
TREM2^+^ and MARCO^+^ macrophage subtypes accumulate in large versus small prostates. Myeloid cell subclustering was conducted by taking original myeloid clusters from the CD45+ scRNA-seq analysis, followed by removal of cells expressing keratin genes or T/B cell specific genes. **a)** UMAP plot indicating 14 myeloid cell subclusters with putative identity. **b)** Violin plots of genes used to identify myeloid subpopulations. **c)** Graph representing the mean +/− SEM of the percentage of cells in each myeloid subcluster among total myeloid cells between samples in large (pink) versus small (blue) prostates (n=10/group). Asterisks indicate significantly changing populations in large versus small prostate tissues by permutation test (*FDR<0.05). Black rectangles indicate clusters with significant upregulation as a percentage of total myeloid cells in large versus small prostates. **d)** UMAP plot of myeloid subclustering from the 20 TZ tissues in (a), colored to highlight cells from small (blue) or large (pink) prostates. Mϕs = macrophages

### BPH macrophages that do not reflect either an M1 or M2 phenotype specifically accumulate with increased prostate volume

Macrophage polarization to M1 and M2 phenotypes has been used widely to suggest either pro-or anti-inflammatory function in various disease states^11^, however, these definitions scarcely reflect macrophage phenotypes in vivo. Alignment of BPH myeloid subpopulations with M1 and M2 transcriptional profiles determined that some clusters have primarily M1-like (subclusters 4, 7, 9, 11-13) or M2-like (subclusters 6 and 10) phenotypes, but other clusters have either a mixed phenotype (subclusters 2 and 8), or do not match well with either polarization profile (subclusters 0, 1, 3, and 5) in tissues (Figure 3a-b, Supplementary Table S1). The cDC2 subclusters were the only subpopulations with M2-like polarization, while macrophage subpopulations aligned with either M1-like, mixed, or neither M1 nor M2 phenotypes (Figures 2a, 3b). These data demonstrate the diversity of BPH macrophages and suggest that *TREM2*^+^ and *MARCO*^+^ macrophages (subclusters 0 and 5, respectively), which increase in relative abundance as the prostate expands, do not fit the spectrum of M1/M2 polarization (Figures 2, 3b). Furthermore, macrophage subpopulations within the neither M1-like nor M2-like category significantly increase as a proportion of total myeloid cells (p<0.0001 via unpaired two-tailed t-test), while all other cluster phenotypes (M1-like, M2-like, and mixed) significantly decrease (p=0.0002, p=0.0472, and p=0.0201 via unpaired t-test, respectively) in large versus small prostate tissues (Figure 3c). To determine whether BPH myeloid cells align with previously defined subpopulations, our myeloid subpopulations were evaluated based on the profiles of prostatic immune cells by Tuong et. al^13^. This evaluation suggests strong alignment, with DC, macrophage, and monocyte clusters overlapping accordingly (Supplementary Figure S4e). Furthermore, BPH macrophage subpopulations do align with synovial macrophage subtypes previously identified by Zhang et. al in rheumatoid arthritis patients^36^ (Supplementary Figure S4f), indicating a transcriptional similarity between myeloid cells in BPH and a defined autoimmune disease.

**Figure 3.**
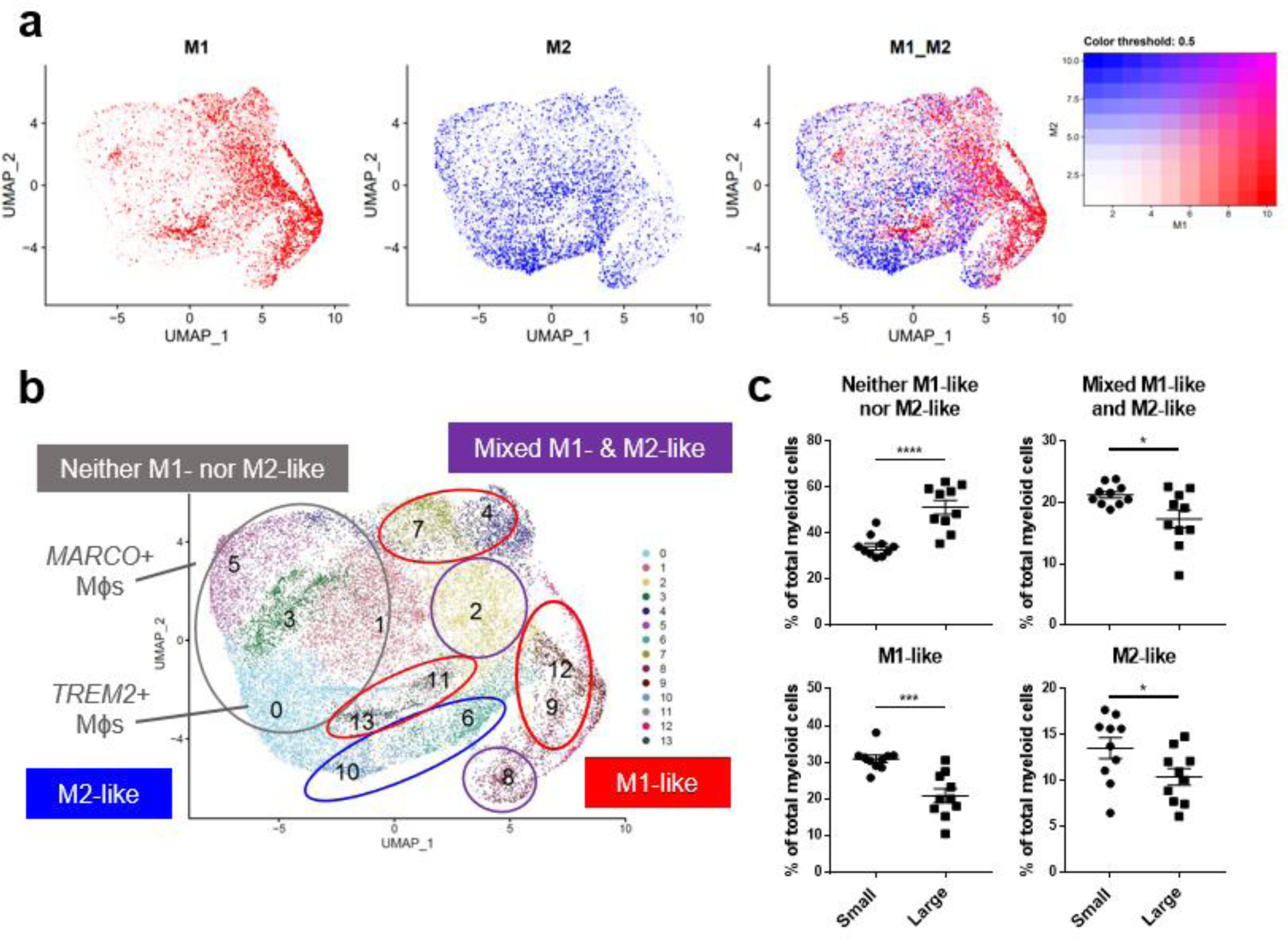
BPH macrophages that do not reflect either an M1 or M2 phenotype specifically accumulate with increased prostate volume. Macrophage (Mϕ) polarization signatures were determined by aligning defined M1 and M2 signature gene sets with the differentially expressed genes for all myeloid cell subclusters. **a)** UMAP plots highlighting individual cells with M1 (red), M2 (blue), or mixed M1 and M2 (purple) signatures, based on the gene module score. More intense color indicates cells with higher scores. **b)** Summary of approximate classification of each myeloid subcluster, based on evaluation of the polarization signature results. Subpopulations are circled with the signature designation, such as M1-like (red), M2-like (blue), mixed M1- and M2-like (purple), and neither M1-nor M2-like (gray). **c)** Graphs indicating the percentage of total myeloid cells from the subclustering analysis within each polarization category, comparing large versus small groups.

### Velocity analysis suggests that TREM2^+^ and MARCO^+^ macrophages are an initial cell state

Given the known plasticity of macrophages, velocity analysis was conducted to determine the trajectory of these subpopulations related to one another. This analysis reveals that the majority of trajectories originate with *TREM2*^+^ macrophages (cluster 0) and *MARCO*^+^ macrophages (cluster 5), while the DCs originate with cluster 10, and clusters 8 and 12 may be a source of monocytes (Figure 4a). While these paths are generally similar between macrophages from both small and large tissue samples (Supplementary Figure S4g), the velocity length in clusters 0 and 5 decrease as prostate size increases, suggesting cells within these clusters may be more stable in that phenotype (Figure 4b)^37^. If less transition from *TREM2*^+^ monocyte-derived or *MARCO*^+^ tissue-resident macrophages to other clusters occurs, this increased stability could be a reason that these specific cell states accumulate in large BPH tissues (Figures 2-4).

**Figure 4.**
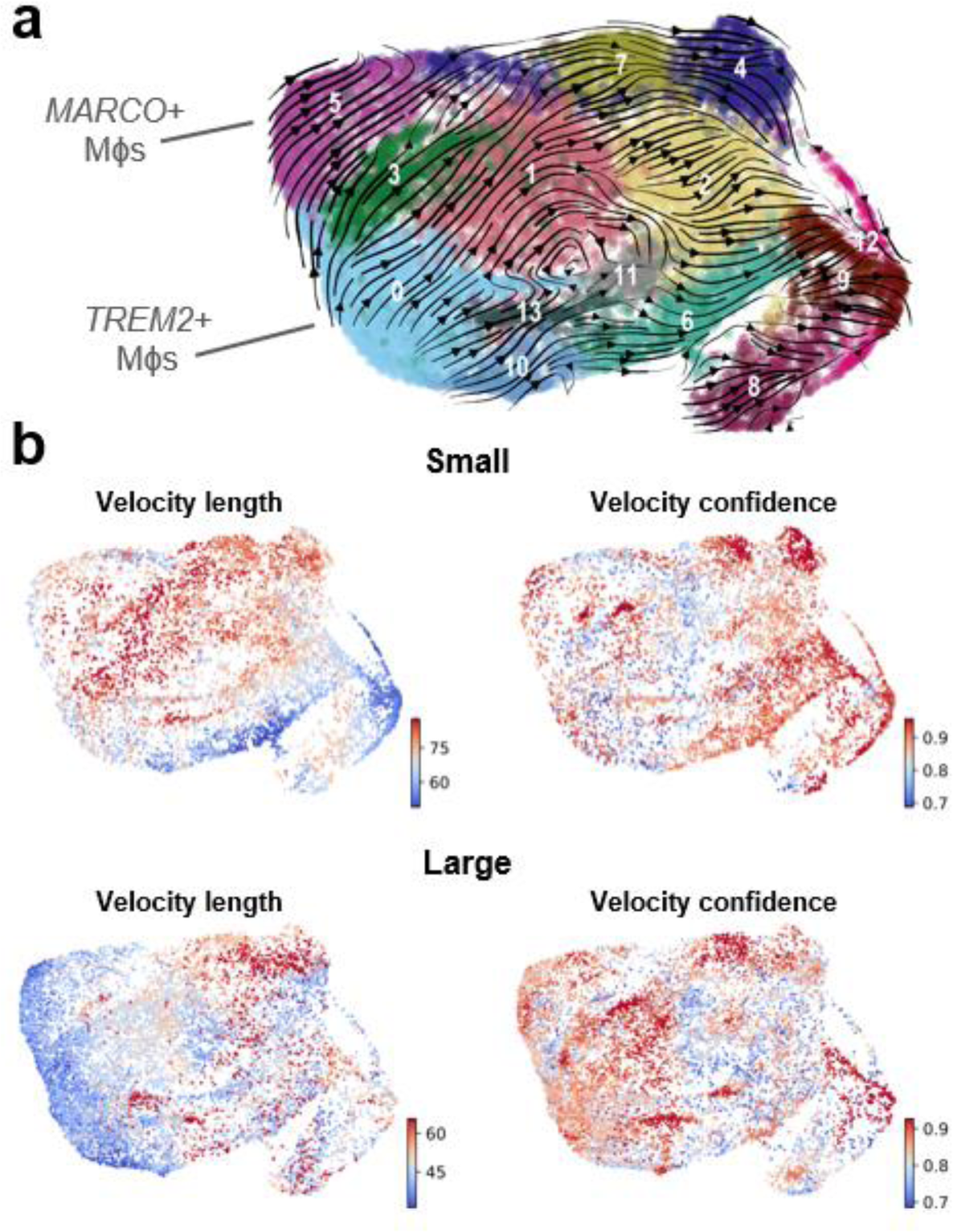
Velocity analysis suggests that TREM2^+^ and MARCO^+^ macrophages are an initial cell state. RNA velocity analysis was conducted to evaluate cellular trajectories within the macrophage (Mϕ) subclustering analysis from small and large groups. **a)** The dynamical model of the velocity stream from n=20 patients was projected onto the UMAP plot generated in Seurat. **b)** Plots indicating velocity length and confidence for the myeloid cells from small or large samples (n=10/group). Red indicates longer velocity length and higher confidence, whereas blue indicates shorter velocity length and lower confidence, where velocity length is proportional to the predicted rate of transition from one cell state to another.

### TREM2^+^ and MARCO^+^ macrophages accumulate intracellular lipid and are associated with clinical characteristics

Since *TREM2*^+^ macrophages accumulate in BPH tissues, immunofluorescence was conducted to determine the localization of these cells in human prostate TZ. TREM2^high^CD68^+^ macrophages are primarily located within stromal areas, while TREM2^low^CD68^+^ macrophages are located both within the stroma and adjacent to epithelial cells (Figure 5a). Evaluating overlapping upregulated marker genes (LogFC>0.5) between *TREM2*^+^ and *MARCO*^+^ macrophages highlighted CD74 as a potential extracellular marker and indicated upregulation of lipid-related genes such as *LIPA*, *APOE*, and *MSR1*. Ingenuity Pathway Analysis identified upregulation of the PPAR signaling pathway in both clusters (Supplementary Figures S5-S6). Furthermore, linear regression of the percent cells in each cluster among total myeloid cells from each patient yielded a significant positive correlation with patient BMI (p=0.0005 and p=0.0009 for TREM2^+^ and MARCO^+^ macrophages, respectively) and IPSS (p=0.0063 and p=0.0008 for TREM2^+^ and MARCO^+^ macrophages, respectively) in this 20-patient cohort (Figure 5b-e). To determine if these cells have more intracellular lipid compared to other macrophages in the prostate, fresh TZ tissues were digested and stained with BODIPY to evaluate neutral lipid levels in the different cell types. HLADR^+^TREM2^+^ macrophages had significantly higher median fluorescent intensity (MFI) for BODIPY than HLADR^+^TREM2^−^ macrophages (p=0.0254 via paired, two-tailed t-test; Figure 5f-g). Using shared marker CD74 for evaluation of intracellular lipid in both TREM2^+^ and MARCO^+^ macrophages determined that HLADR^+^CD74^+^ macrophages have higher BODIPY MFI than HLADR^+^CD74^−^ macrophages (p=0.0531 via paired, two-tailed t-test; Supplementary Figure S7). The gating strategy for macrophage subset identification is indicated in Supplementary Figure S8. These studies demonstrate that accumulating macrophages in BPH tissues increase intracellular neutral lipid stores and positively correlate with patient symptoms.

**Figure 5.**
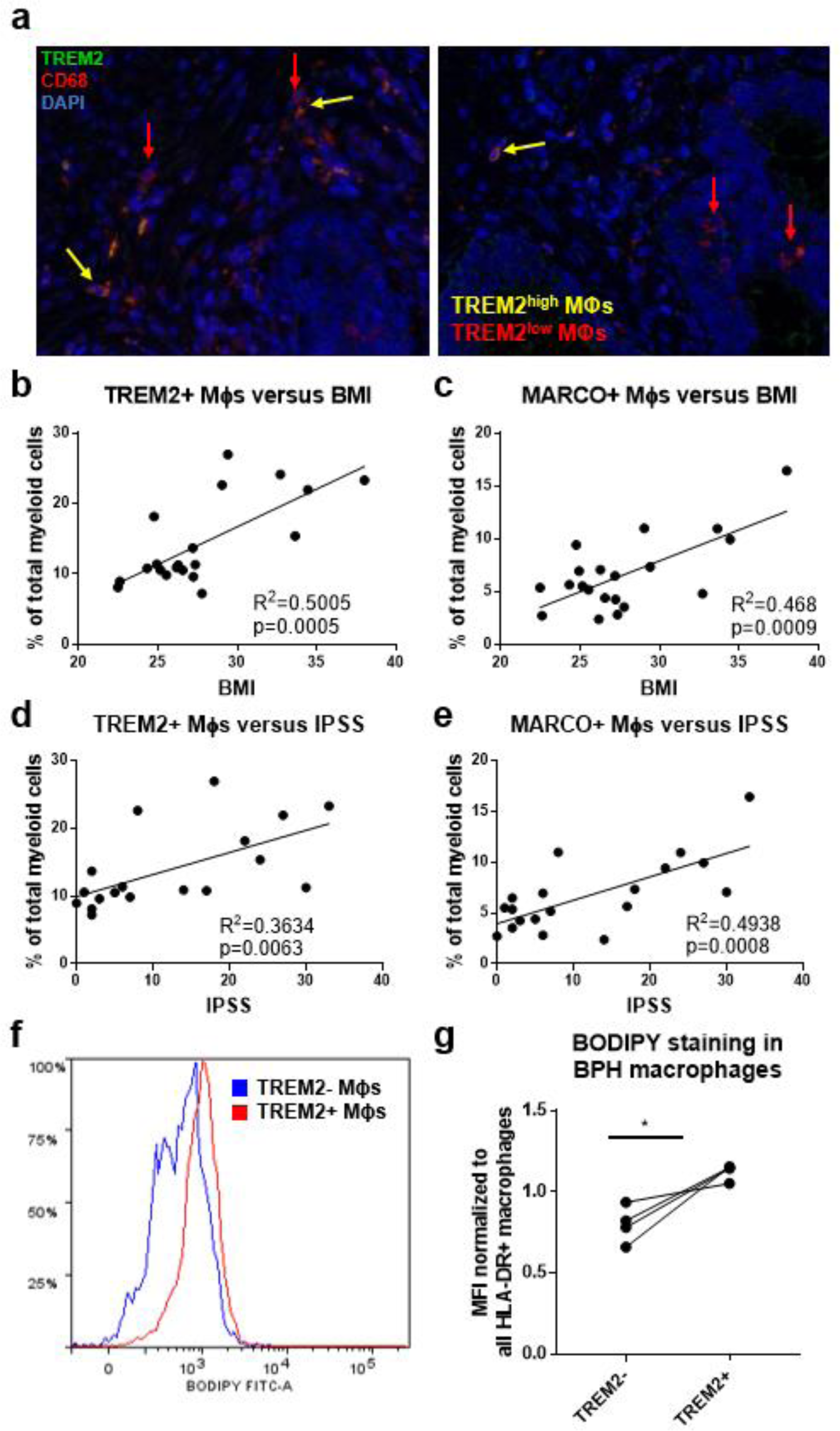
TREM2^+^ and MARCO^+^ macrophages are associated with IPSS and BMI and accumulate intracellular lipid. **a)** Immunofluorescence image of human BPH tissue indicating TREM2 (green) co-localization with macrophage (Mϕ) marker CD68 (red). DAPI (blue) indicates nuclei. Red arrows identify TREM2^low^ macrophages and yellow arrows indicate TREM2^high^ macrophages in the prostate stroma. **b-e)** Linear regression of the percentage of cells in TREM2+ cluster 0 (b, d) or MARCO+ cluster 5 (c, e) versus patient BMI (b-c) or IPSS (d-e). **f-g)** Human BPH tissues were digested and stained for flow cytometry analysis. BODIPY staining intensity was measured in TREM2^+^ versus TREM2^−^ subpopulations. **f)** Example histogram of BODIPY median fluorescence intensity (MFI) in TREM2^+^ (blue) versus TREM2^−^ (red) macrophage subpopulations. **g)** Quantitation of (f) from 2 independent experiments, where the BODIPY MFI was normalized to the MFI for all HLA-DR+ macrophages in each sample.

### T cell subclustering indicates no accumulation of T cell subpopulations in large versus small volume prostates

Since lipid-rich myeloid cells have been shown to have decreased antigen presentation capability resulting in diminished T cell activation^17^, we evaluated whether T cell subpopulations change as prostate size increases. Subclustering analysis of CD3^+^ T cells and CD3^+^ NK cells from all 20 samples was performed and visualized to produce 14 transcriptomically-distinct subpopulations (Supplementary Figure S9a). Identification of subclusters was determined empirically with both CD4 and CD8 cell surface protein expression and with differentially expressed genes (DEGs) between clusters (Supplementary Figures S9-S10)^38–41^. There are no significant differences in T cell subclusters due to individual patient samples (Supplementary Figure S11a). When comparing cells from large versus small tissues, only CD16^−^ NK cells demonstrate an increase in relative proportion among total CD45^+^ cells, while the majority of T cell subclusters are unchanged with the exception of a decrease in the relative proportion of CD8^+^ T effector/central memory (T_EM/CM_) and metallothionein expressing T (MT-T) cells (Supplementary Figures S10, S11b-c). Expression of metallothionein genes have also been used as markers of T cell exhaustion^42^. While it is important to note that CD45^+^ inflammatory cells in general accumulate as prostate size increases^6,43^, there is no significant increase in the relative abundance of T cell subclusters that associate with increasing prostate volume.

### Lipid-loaded THP-1 macrophages increase prostatic epithelial and stromal cell proliferation

The neutral lipid composition of lipid droplets is primarily triacylglycerols or cholesterol esters. To examine the impact of excess lipid on the microenvironment, THP-1 monocyte-derived macrophages were pre-loaded with either cholesterol or oleic acid (OA) prior to transwell co-culture with prostatic epithelial (NHPrE-1) or stromal (BHPrS-1) cell lines. Both epithelial and stromal cells demonstrate significantly increased proliferation when co-cultured with OA-loaded macrophages (p_adj_=0.0113 and p_adj_=0.0138 via one-way ANOVA, respectively), but not cholesterol-loaded macrophages, compared to control macrophages (Figure 6a-b). To visualize lipid-loading in a subset of macrophages, BPH patient tissues were stained with both Oil Red O and CD68 via IHC. This analysis demonstrates the presence of both lipid-rich and lipid-poor macrophages in the prostate stroma of BPH patients (Figure 6c). These data support the idea that specific BPH macrophage subpopulations accumulate intracellular lipid and directly stimulate increased epithelial and stromal cell proliferation, resulting in increased prostatic hyperplasia and patient symptoms.

**Figure 6.**
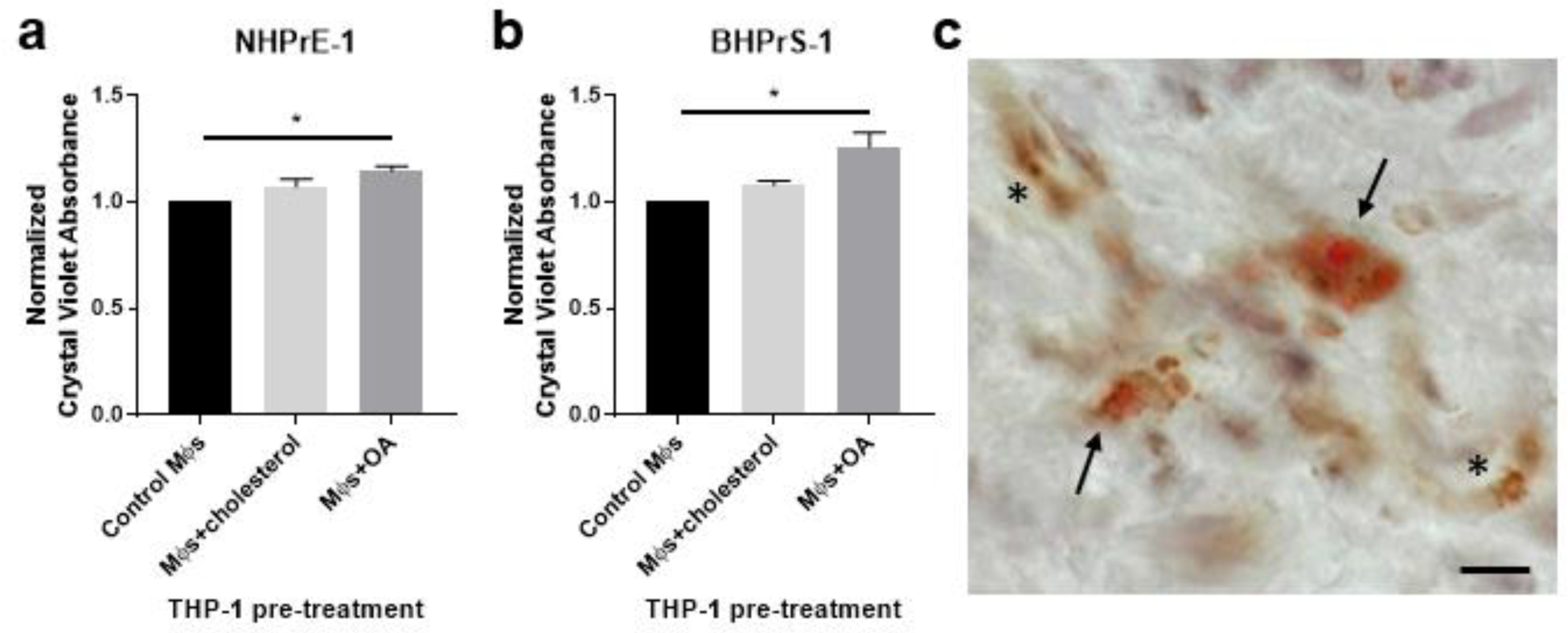
Lipid-loaded THP-1 macrophages increase prostatic epithelial and stromal cell proliferation. THP-1 monocytes were differentiated to macrophages (Mϕs) by 48-hour treatment with PMA. Cells were lipid-loaded with either cholesterol or oleic acid (OA) as indicated. **a-b)** Crystal violet growth assay representing cell proliferation of NHPrE-1 epithelial cells (a) and BHPrS-1 fibroblasts (b) after 4 days of transwell co-culture with control, cholesterol-treated, or OA-treated macrophages. Graphs represent the mean +/− SEM of the average values from three independent experiments, normalized to the cells incubated with control macrophages. **c)** Human BPH tissues were stained for neutral lipids with Oil Red O (red) and macrophage marker CD68 (brown) via IHC. Image demonstrates lipid-rich macrophages (arrows) and lipid-poor macrophages (asterisks) in the prostate stroma. Staining was completed using two BPH patient tissues, unique from the single-cell patient cohort. Scale bar = 20 µm.

## Discussion

The results from these studies provide a detailed cellular description of leukocytes within the prostate transition zone and demonstrate the specific accumulation of TREM2^+^ and MARCO^+^ macrophages as prostate size increases. The enriched macrophage subpopulations have elevated intracellular lipid and altered PPARγ signaling compared to other BPH macrophages. Since these lipid-rich macrophages positively correlate with urinary symptom scores (IPSS) and lipid-loaded macrophages stimulate epithelial and stromal proliferation in vitro, the present data support that TREM2^+^ and MARCO^+^ macrophages promote prostatic hyperplasia and associated urinary symptoms in BPH patients.

Data herein highlight the complexity of macrophage phenotype classification. Macrophage plasticity is widely discussed and could be the result of ontogeny as well as responses to stimuli in the microenvironment. This highlights a need to understand macrophage cell states in relation to organ and disease status, so that macrophage function can be characterized in a targeted manner. It is notable that the transcriptional phenotypes of resident macrophages can differ between organs even though function remains the same^44^. Transcriptional profiling of macrophage cell states is further complicated by technical differences in processing and sequencing between studies within the same organ^13,14,45^, highlighting a need for standardization of methodology. Nonetheless, the successful alignment of BPH macrophage subpopulations to those of synovial macrophages in rheumatoid arthritis illustrates the association between inflammation in BPH and autoimmune disease^6,36^ and confirms that inflammatory commonalities between conditions exist. The blend of macrophage phenotypes observed in BPH is suggestive of a chronic, non-resolving inflammatory state. Further studies are needed to investigate the involvement of T cell activation and function in the chronic inflammatory process in BPH tissues, as this will determine whether BPH categorizes into either an autoimmune or an autoinflammatory disease process.

BPH macrophage phenotypes described in these studies also indicate a need to update macrophage polarization categories. Previous studies have demonstrated that cells resembling M1/M2 polarization are spread among macrophage subclusters with varying transcriptional profiles^45,46^. Considering the presence of a progressive inflammatory state in BPH tissues as prostate size increases, it is reasonable to predict that macrophages with an M1 phenotype would accumulate in large versus small prostates. However, our data indicate that subpopulations of macrophages expressing neither M1-nor M2-like polarization signatures preferentially accumulate as prostate size increases, while M1-like macrophages actually *decrease* in relative abundance in large versus small prostates. Activation of PPARγ in lipid-loaded macrophages may be involved in the polarization signature since this transcription factor is known to inhibit expression of some inflammatory genes^18^. PPARγ signaling may also be responsible for the high expression of *MRC1* (CD206), a marker often used to indicate M2 polarization, in MARCO^+^ macrophages^47^.

Accumulation of macrophage subpopulations in BPH tissues warrants the investigation of prostatic macrophage origin. Lineage tracing would be necessary to verify whether the macrophage subpopulations identified in these studies are derived from yolk-sac macrophages, fetal monocytes, or bone marrow hematopoietic stem cells^48,49^. However, analysis of previously defined conserved tissue-resident macrophage markers in this work suggests that some BPH macrophages are tissue-resident^28,29^. The analysis of cell cycle genes provides evidence that TREM2^+^ and MARCO^+^ macrophages are proliferating, but it is also possible that these cells are infiltrating the tissue from circulating monocytes given their stromal localization. Indeed, tissue-resident macrophages can be derived from monocytes as well as the yolk-sac or fetal liver^50^. The velocity analysis also indicates these cells become more stable as prostate size increases, which is perhaps due to a biological function that keeps these cells from transitioning to other transcriptional phenotypes.

While lipid metabolism in prostate cancer has attracted significant attention, there has been minimal investigation of these pathways in BPH tissues. Recent work by Popovics, et. al has indicated the presence of “foamy” macrophages in the glandular lumen of BPH tissues^51^, but through our studies we discovered that a subpopulation of lipid-rich macrophages additionally accumulate in the stroma of human prostates and may contribute to cell proliferation and LUTS. Work in the steroid hormone imbalance model of BPH also suggests that TREM2^hi^ macrophages exist in the mouse prostate after hormone treatment^52^.

The mechanism by which TREM2^+^ or MARCO^+^ macrophages accumulate lipid is unclear. If lipid-rich macrophages in BPH are indeed a combination of tissue-resident (MARCO^+^) and monocyte-derived (TREM2^+^) cells as the cluster markers suggest, it indicates that both systemic and local variables could contribute to lipid-loading. Previous studies have determined that high fat diet can increase macrophage infiltration and alter polarization status in a prostate cancer model and demonstrated that lipid-loaded tumor-associated macrophages (*MARCO*^+^) support prostate tumor growth^14,53^. In BPH, positive correlations of *TREM2*^+^/*MARCO*^+^ macrophage abundance with patient BMI also supports a model where elevated systemic lipids serve as a source of lipid uptake, while the TREM2 or MARCO proteins themselves could serve as lipid receptors^14,19^. Intracellular reprogramming of lipid metabolism regulators or pathways could also contribute to lipid accumulation^54^. Further studies are needed to explore both the cause(s) and downstream effects of the net gain of neutral lipid in BPH macrophages.

The scRNA-seq data presented in these studies have limitations. The classification of macrophage subpopulations was primarily empirical due to the lack of defined cell states as discussed above. However, the focus on immune cells in this study compared to previous datasets^55–57^ provides both more cells and more genes/cell for immune subcluster identification. The presence of lipid-rich macrophages was also verified in distinct BPH tissues. All identified immune subpopulations will be further investigated for unique biological functions in future studies. Although the scRNA-seq study was limited to 20 total patients and primarily examines the changes related to excessive prostate size in BPH, significant changes within CD45+ leukocytes, specifically among macrophages, were identified in large versus small prostates. Finally, while the *in vitro* studies were limited to the use of THP-1 cells, which do not reflect macrophages in all disease states, the data support a model in which lipid-loading in macrophages promotes prostate cell proliferation. It is not known which lipids are most relevant to the inflamed BPH tissue microenvironment, so the naturally abundant fatty acid, oleic acid, was used for lipid-loading and downstream proliferation studies.

Although these data indicate that TREM2^+^ macrophages positively correlate with patient-reported urinary symptoms via the IPSS, the question remains whether these cells drive BPH progression and subsequently worsen symptoms or are accumulating as a response, such as in an effort to resolve chronic inflammation. TREM2^+^ cells have been shown to have beneficial properties in hepatic and renal tissue damage, but blockade of these cells restores anti-tumor immunity^58–61^. There is evidence for both pro- and anti-atherosclerotic properties of TREM2^33,62^, and it remains possible that lipid-loaded macrophages have a role in restraining inflammation via PPAR^63^. In the tumor microenvironment, lipid-loading in tumor-associated macrophages decreases phagocytosis and increases PD-L1 expression, resulting in an immunosuppressive environment for tumor growth^17^. However, increased TREM2 expression in myeloid cells can increase phagocytic ability and is significantly elevated in patients with inflammatory bowel disease^19,21^. Thus, the function of TREM2^+^ cells may differ based on organ/disease context, with evidence that these cells promote progression of some autoimmune diseases.

These studies suggest potential therapeutic targets not previously investigated in BPH. Therapeutic targeting of inflammation in BPH patients using TNF-antagonists is underway (NCT06062875, clinicaltrials.gov). TNF blockade reduces macrophage accumulation in human and mouse prostate tissues^6^, although the macrophage subtypes involved have not yet been determined. Lipid metabolism pathways have been investigated as therapeutic targets in prostate cancer patients, but not in BPH patients. Data here suggests that infiltrating lipid-rich macrophages may be a more specific inflammatory cell target in BPH patients. Further studies characterizing the role of these cells in BPH will yield new therapeutic strategies, such as lipid-based or cell-specific therapeutics, that are necessary to decrease chronic inflammation and voiding symptoms.

## Methods

All research studies were conducted in accordance with applicable local, state, and national regulations. Human studies were conducted under the approval of the Endeavor Health (formerly known as NorthShore University HealthSystem) Institutional Review Board (IRB). No patients received compensation for participation in these studies.

### Isolation of CD45^+^ cells from human tissues

Human prostate transition zone tissues were ethically procured with the IRB-approved NorthShore Urologic Disease Biorepository and Database with informed consent and de-identified clinical annotation. Small prostate tissues were obtained from 10 male patients undergoing robotic-assisted laparoscopic prostatectomy (RALP) for prostate cancer, International Prostate Symptom Score (IPSS) of <15, Gleason 6-7, and estimated prostate volume of <40 grams by imaging with transrectal ultrasound (TRUS), CT scan, or MRI. Tissues were also obtained from 10 male patients with large volume prostates (estimated prostate size of >90 grams) who were undergoing either RALP for prostate cancer (Gleason 6-7) or simple prostatectomy for BPH. Clinical information for the 20 patients is included in Supplementary Data S1. The transition zone (TZ) was dissected similarly for both small and large prostate tissues and were pathologically verified to have no (or minimal) cancer burden. Tissue pieces were separated for formalin-fixed paraffin-embedded (FFPE) histology or digested and prepared for fluorescence activated cell sorting (FACS). Tissues were minced, then digested while shaking at 37°C for 2 hours in 200 U/mL Collagenase I (Gibco) + 1 mg/mL DNase I (Roche) + 1% antibiotic/antimycotic solution in Hank’s Balanced Salt Solution. Digestion solution was replaced with TrypLE Express dissociation reagent (Gibco) and allowed to shake at 37°C for 5-10 minutes. Digested samples were neutralized in RPMI + 10% FBS, then mechanically disrupted by pipetting repeatedly. Samples were passed through a 100µm cell strainer, then washed. Red blood cells were lysed in a hypotonic buffer before the cell suspension was stained with Zombie Violet (Biolegend) and blocked with Human TruStain FcX blocking antibody (Biolegend). CD45-PE [clone HI30], EpCAM-APC [clone 9C4], and CD200-PE/Cy7 [clone OX-104] antibodies (Biolegend) were added to stain samples in preparation for FACS on a BD FACSAria II. Approximately 100,000 viable CD45^+^CD200^−^EpCAM^−^ cells were sorted for downstream analysis.

A separate tube of the digested cell suspension was labeled for flow cytometry analysis of immune cells and stained with Zombie Violet (Biolegend) as well as CD45-FITC [clone HI30], CD11b-PE/Cy7 [clone ICRF44], CD19-APC/Cy7 [clone HIB19], CD3-APC [clone UCHT1], CD4-PE [clone RPA-T4], and CD8-BV510 [clone RPA-T8] antibodies. Information for all antibodies can be found in Supplementary Table 2.

### scRNA-seq of CD45^+^ cells

FACS-isolated cells were spun down and washed at least twice before loading onto the 10X Chromium System (10X Genomics), with Single Cell 3’ Library & Gel Bead Kit, v3.0 reagents. Cells from three small and nine large tissues were stained with TotalSeq-B Antibodies (Biolegend) for CITE-seq analysis. Antibodies for CD3 [clone UCHT1], CD4 [clone RPA-T4], CD8 [clone RPA-T8], CD19 [clone HIB19], CD11b [clone ICRF44], and CD56 (NCAM) [clone 5.1H11] were used following the manufacturer’s instructions before loading into the Chromium System. Cells were loaded for downstream evaluation of 5,000 cells/sample and cDNA amplification and library preparation were conducted according to the manufacturer’s instructions. Libraries were sent to the Purdue Genomics Core Facility for post-library construction quality control, quantification, and sequencing. A high-sensitivity DNA chip was run on an Agilent Bioanalyzer (Agilent) per the recommendation of 10x Genomics. Additional quality control was performed by running a denatured DNA pico chip (Agilent) followed by an AMPure cleanup (Beckman Coulter). Final library quantification was completed using a Kapa kit (Roche KK4824) prior to sequencing. Normalized pools were sequenced using a NovaSeq S4 flow cell on a NovaSeq 6000 system (Illumina) with 2*x*150 base-pair reads at a depth of 50,000 reads/cell. Libraries generated from cell surface protein labeling were sequenced at a depth of 5,000 reads/cell.

### Initial processing and quality control of scRNA-seq data

Sequencing reads were processed using the CellRanger pipeline v3.0.0 (10x Genomics). Specifically, CellRanger mkfastq was run to generate FASTQ files using the flag “—use-bases-mask=Y26n*,I8n*,n*,Y98n”, ignoring dual indices, and allowing 0 mismatches. Alignment to the ENSEMBL GrCh38 human reference genome, barcoded filtering, and counting unique molecular identifiers (UMIs) were performed using the program CellRanger count. R version 4.1.2 and Bioconductor version 3.8 were used in scRNA-seq statistical analyses unless explicitly mentioned. Cells with between 1,000 and 10,000 observed genes were retained, and less than 22% of all reads were mapped to mitochondrial genes. Run metrics and a summary of the data produced by the scRNA-seq analyses are shown in Supplementary Table S3. Raw and processed scRNA-seq data were uploaded to GEO and are available under accession numbers GSExxxxxx.

### Clustering and downstream analysis of combined leukocyte scRNA-seq data

Data normalization and unsupervised clustering of scRNA-seq data were performed using Seurat version 4.1.0^64–68^. The scTransform package v.0.3.3^69^ was used to normalize and scale the data, ultimately removing unwanted heterogeneity by regressing the UMI counts, cell cycle scores, and the percent of reads mapping to mitochondrial genes.

The first 15 principal components of the scaled data were then used for downstream analyses. Unsupervised clustering was performed using graph-based approaches to construct K-nearest neighbor graphs (K=15) in Seurat. The Seurat implementation of the Louvain method for community detection was then used to identify clusters of similar cells by optimizing the modularity function^70^. The R package clustree v 0.4.4^71^ was used to select the optimal resolution of 0.5, ultimately allowing the selection of a resolution that provides stable, resolved clusters. Marker genes were identified for each cluster using the Wilcoxon rank sum test^72^, as implemented within Seurat. These markers were considered statistically significant at a 1% false discovery rate (FDR). Differentially expressed genes with FDR<5% were identified between small and large sample groups using the Wilcoxon rank sum test after “pseudobulking” across cells within samples using the function AggregateExpression() in Seurat. The Benjamini-Hochberg method^73^ was used to correct P-values for multiple testing. A permutation test was used to identify clusters of cells that significantly increase or decrease in number between large and small samples, as described in^74^. Gene ontology terms, KEGG pathways, and Reactome pathways that are enriched amongst the cluster marker genes and the differentially expressed genes were identified by the package ClusterProfiler v.4.2.2^75,76^ using all of the detected genes from the dataset as the background and controlling FDR at 5%.

### Subclustering and identification of macrophage cell states

For the macrophage clustering, clusters 2, 5, 9, 11, 12, and 13 were selected from the combined clustering, re-normalized, integrated using anchor-based clustering, and communitites were detected. Next, cells expressing *CD3D*, *CD3E*, *CD3G*, *CD19*, *CD20*, and *CD79A* were removed. Cells expressing *CD3* genes were moved to the T cell subclustering analysis. Finally, cells expressing keratins (*KRT5*, *KRT8*, *KRT15*, *KRT18*, *KRT19*, *KRT81*, or *KRT86*) were removed. Subsequently, the remaining cells were re-normalized, clustered, and communities detected. The top 12 principal components were used in clustering the cells and an optimal resolution of 0.4 was used for community detection. Identification of marker genes, differentially expressed genes, and enriched pathways were performed as described above for the combined leukocyte analysis.

The BPH macrophage subsets were annotated using macrophage subsets from the synovial tissue of patients diagnosed with rheumatoid arthritis^36^. Garnett^30^ was used to train a classifier using default parameters and the top 20 statistically significant upregulated marker genes from each subset, ranked by fold-change as features.

### Macrophage polarization signature analysis

An analysis was performed to quantify macrophage polarization, specifically identifying cells with enrichment of M1 (LPS+IFNg treatment) or M2 (IL-4, IL-13 treatment) gene signatures. The gene signatures from Becker, et. al^11^ were used, where the M1 signature included *ADAM28*, *AIM2*, *ANKRD22*, *APOBEC3A*, *APOL1*, *APOL3*, *BATF2*, *C1R*, *C1S*, *CCL19*, *CD38*, *CD40*, *CD80*, *CFB*, *CLEC4D*, *CXCL10*, *CXCL9*, *CYBB*, *DUCP10*, *DUSP6*, *ETV7*, *FAM49A*, *FAM65B*, *FCGR1B*, *FPR2*, *GADD45G*, *GBP1*, *GBP2*, *GBP4*, *GBP5*, *GCH1*, *GK*, *GPR84*, *GUCY1A3*, *HERC5*, *HESX1*, *HLA-F*, *IFI27*, *IFI35*, *IFI44L*, *IFIH1*, *IFIT2*, *IFIT3*, *IFITM1*, *IFITM2*, *IL15*, *IL15RA*, *IL32*, *INHBA*, *IRF1*, *IRF7*, *ISG15*, *ISG20*, *ITGAL*, *ITGB7*, *LAG3*, *LAMP3*, *LIMK2*, *LRRK2*, *MUC1*, *MX1*, *NAMPT*, *NFKBIZ*, *OAS1*, *OAS2*, *OAS3*, *OASL*, *OPTN*, *PAG1*, *PARP14*, *PCNX*, *PDE4B*, *PIM1*, *PRKAR2B*, *PSMB9*, *PTGS2*, *RARRES3*, *RCN1*, *RHBDF2*, *RSAD2*, *SAT1*, *SCO2*, *SEPT4*, *SERPING1*, *SLAMF7*, *SLC22A15*, *SLC25A28*, *SLC31A2*, *SLC6A12*, *SLC7A5*, *SNTB1*, *SNX10*, *SOCS3*, *SOD2*, *STAT1*, *STAT3*, *STX11*, *TAP1*, *TNFAIP6*, *TNFSF10*, *TRIM69*, *UBE2L6*, *USP18*, *VAMP5*, *WARS*, and *XRN1*, while the M2 signature included *ADAM19*, *ALOX15*, *ARRB1*, *BZW2*, *CARD9*, *CCL13*, *CCL17*, *CCL23*, *CD1A*, *CD1C*, *CD1E*, *CDR2L*, *CHN2*, *CLEC4A*, *CLIC2*, *CMTM8*, *CRIP1*, *CTSC*, *DUSP22*, *EMILIN2*, *ESPNL*, *F13A1*, *FOXQ1*, *FSCN1*, *FZD2*, *GALNTL4*, *GATM*, *GPD1L*, *GSTP1*, *ITM2C*, *KCNK6*, *MAOA*, *MAP4K1*, *MAPKAPK3*, *MFNG*, *MS4A6A*, *NMNAT3*, *OSBPL7*, *P2RY11*, *PALLD*, *PAQR4*, *PELP1*, *PLAU*, *PON2*, *PPP1R14A*, *PTGS1*, *RAMP1*, *REPS2*, *RGS18*, *RRS1*, *S100A4*, *SEC14L5*, *SHPK*, *SPINT2*, *TGFB1*, *TMEM97*, *VCL*, *SNF789*. Control features were selected using binning based on average expression, and control features were randomly selected from each bin, as implemented in Seurat^70^. We calculated an aggregate M1 and M2 gene module score based on the average expression levels of each gene on a single cell level, subtracting out the aggregated expression of control feature sets. We confirmed the gene-module-based classification of clusters using the M1 and M2 signature genes mentioned above to perform a gene set enrichment analysis in QuSAGE v.2.28.0^77^. Classifications to either M1-like, M2-like, Mixed M1-& M2-like, or Neither M1-nor M2-like were made based on log(fold-change) and p<0.01. The QuSAGE results are indicated in Supplementary Table S1.

### Subclustering and identification of T cell subsets

T cells and CD3+ NK cells were then subset out of this combined clustering by taking clusters 0, 1, 3, 4, 7, 10, 16, and 17 and combined with cells from all other clusters that express *CD3D*, *CD3E*, or *CD3G*. Next, cells expressing *CD68*, *CD19*, *CD20*, and *CD79A* were removed. Cells expressing *CD68* were moved to the macrophage subclustering analysis. Finally, cells expressing keratins (*KRT5*, *KRT8*, *KRT15*, *KRT18*, *KRT19*, *KRT81*, or *KRT86*) were removed. Data was re-normalized, and integrated using anchor-based clustering, and communities were detected. The top 13 principal components were used in clustering the cells and a resolution of 0.5 was used for community detection. Identification of marker genes, differentially expressed genes, and enriched pathways were performed as described above for the combined leukocyte analysis.

### RNA velocity analyses

RNA velocity analyses were performed on the macrophage subclusters. Loom files were created from CellRanger output using Velocyto^78^ v.0.17.17. RNA velocity analyses were performed using the embeddings and clusters identified by Seurat after exporting the genes, counts, metadata, and embeddings from the macrophage and T cell subclustering results. An RNA velocity analysis was performed using scVelo^79^ v. 2.4.0, employing the dynamical model used for velocity analysis. Following scVelo analysis, CellRank^80^ v. 1.5.1 was run in an attempt to uncover cellular dynamics. The scVelo and CellRank analyses were applied to the integrated macrophage subclustering dataset.

### Immunofluorescence

Human FFPE BPH tissue sections of 5µm thickness were mounted on slides and prepared for immunofluorescence (IF). Sections were deparaffinized in xylene, treated with hydrogen peroxide (H_2_O_2_) for endogenous peroxidase removal, and rehydrated using gradient ethanol concentrations. Heat-based antigen retrieval was completed with Antigen Unmasking Solution (Vector H-3300) and blocking was conducted using 10% goat serum in 1% BSA solution. Primary antibodies targeting TREM2 (Thermo 702886) and CD68 (Dako M0814) were incubated overnight at 4°C, followed by secondary antibody labeling with anti-mouse AF594 (Invitrogen A11032) and anti-rabbit AF488 (ThermoFisher A27034) before mounting with DAPI.

### BODIPY staining of macrophage subpopulations from human prostate tissue

Prostate TZ tissues were obtained through biobank (as above), minced, and digested in 200 U/mL Collagenase I (Gibco), 1 mg/mL DNAse I (Fisher), and 1% antibiotic/antimyotic solution in Hanks Balanced Salt Solution (Fisher) shaking at 37°C for 2.5 hours. TrypLE Express dissociation reagent (Gibco) was used to dissociate tissues for an additional 5-10 minutes at 37°C. The neutralized cell suspension and remaining tissue pieces were mechanically dissociated using a 16G needle and passed through 100 µm and 70 µm cell strainers. The cell suspension was washed and Ammonium-Calcium-Potassium (ACK) buffer was used to lyse red blood cells. Samples were stained with Zombie Violet (Biolegend 77477) and Human TruStain FcX Blocking Reagent (Biolegend 422302) for 10 minutes. The cells were washed and stained with 2 µM BODIPY 493/503 (ThermoFisher D3922) for 15 minutes at 37°C. After washing, the cells were stained with CD3-BV605 [clone UCHT1], CD19-BV605 [clone HIB19], CD56-BV605 [clone 5.1H11], CD45-PE/Cy7 [clone HI30], HLADR-AF700 [clone L243], CD1c-BV650 [clone L161], CD74-PE [clone LN2] antibodies (Biolegend), and TREM2-APC [clone 237920] before flow cytometry on a BD FACSAria Fusion. The gating strategy for the analysis of macrophages is in Supplementary Figure S8.

### Transwell co-culture and proliferation assay

THP-1 cells were purchased and authenticated from ATCC (STRB0424) and used within 20 passages from testing. THP-1 cells were cultured precisely as indicated by ATCC. NHPrE-1 and BHPrS-1 cell lines were isolated and cultured as benign epithelial and stromal prostatic cell models^81,82^. The authentication of NHPrE-1 (STRA3441) and BHPrS-1 (STRB0418) cells was completed by ATCC, and all experiments were conducted within 20 passages from testing.

THP-1 cells were differentiated with 10 ng/mL phorbol 12-myristate 13-acetate (PMA) (Sigma) in 0.4 µm transparent PET membrane inserts (Falcon) for 48 hours. After differentiation, PMA was removed and the macrophages were treated with or without 100 µM oleic acid (Sigma O3008) or 10 µM cholesterol (Sigma C4951). After 48 hours, the inserts were washed and transferred into wells containing epithelial and stromal cells (NHPrE-1 and BHPrS-1) for co-culture in media containing 1% serum. The co-cultures were incubated for four days, followed by a crystal violet growth assay. Briefly, epithelial or stromal cells were fixed with 4% paraformaldehyde, stained with 0.1% crystal violet, and then solubilized with 10% acetic acid. A spectrophotometer was used to obtain absorbance values at 590 nm.

### Immunohistochemistry and Oil Red O staining

Fresh, frozen tissue sections from two simple prostatectomy patients (independent of the scRNA-seq cohort) were dried onto slides and fixed in 10% formalin. Sections were permeabilized with 0.25% Triton X-100 and treated with 1.5% H_2_O_2_ diluted in PBS to remove endogenous peroxidases. Sections were blocked with 5% horse serum in 1% BSA solution and incubated with the anti-CD68 primary antibody (KP1, Dako) for 60 minutes at room temperature. After washing, secondary antibody staining and ABC labeling were completed using the Vectastain Universal Elite ABC HRP kit (Vector PK-6200). Next, sections underwent Oil Red O staining by pretreating with propylene glycol for 4 minutes, followed by Oil Red O incubation for 1.5 hours while rocking. DAB was completed after washing off the Oil Red O with ddH_2_O. Finally, IHC was completed with hematoxylin counterstain and coverslipping with aqueous mounting medium.

### Statistical Analysis

Statistical significance of *in vitro* assays was completed using a two-way analysis of variance (ANOVA), patient characteristics were compared using a student’s t-test, and immune cell compartments were correlated using linear regression using Prism software version 7.05 (GraphPad). IHC counts were evaluated using a nested t-test (Prism, v8). A p-value of less than 0.05 was considered significant. In data figures, significance is indicated by *=p<0.05, **=p<0.01, ***=p<0.001, and ****=p<0.0001.

## Data Availability

The scRNA-seq data is available in GEO under accession numbers GSExxxxxx. Any further information about tissue resources and reagents associated with these studies should be directed to, and will be fulfilled by, the corresponding author upon reasonable request.

## Code Availability

The R scripts used to perform the scRNA-seq analysis are available at https://github.com/natallah/Lipid-rich_macrophages_accrue_in_large_prostates_2024 and a stable version of the scripts at the time of manuscript submission is available through Zenodo^83^ at https://doi.org/10.5281/zenodo11494628 and is associated with an Apache 2.0 license, allowing users to freely use the scripts for any purpose.

## Acknowledgements

The authors appreciate Phillip SanMiguel and the Purdue University Genomics Facility for their aid in scRNA-seq library normalization and sequencing. The authors are grateful to the NorthShore Biospecimen Repository, pathology assistants Taylor Marvin and Amanda Proulx, clinical research coordinator George Javich, and especially patients who have donated their tissue for research, without which much of this work would not have been possible. We also acknowledge Jeffrey Gaynes for his technical assistance.

## Funding

This work was funded by 1P20DK116185 (S.W.H. and T.L.R.) and R01DK117906 (S.W.H.) from NIDDK, the Purdue University Institute for Cancer Research (NIH grant P30CA023168), the IU Simon Cancer Center (NIH grant P30CA082709), and the NorthShore University HealthSystem Research Institute/Medical Group Pilot Grant (R.E.V.). This work was also generously supported by the Collaborative Core for Cancer Bioinformatics, the Walther Cancer Foundation, and the Rob Brooks Fund for Precision Prostate Cancer Care.

## Author Contributions Statement

S.E.C., T.L.R., S.W.H., O.E.F. and R.E.V. created the study concept; N.A.L., S.E.C., O.E.F., T.L.R., S.W.H., and R.E.V. designed research studies; E.M., P.F., Y.F., P.T., M.A., S.B., A.P.G., A.H., B.T.H., O.E.F., S.W.H., and R.E.V. were involved in acquisition of patient tissues or accessing medical records; N.A.L., H.K., and A.K. performed statistical and bioinformatics analyses; N.A.L., E.M., P.F., Y.F., G.M.C., M.M.B., and R.E.V. performed data acquisition and all authors contributed to data analysis and interpretation. N.A.L., E.M., S.E.C., S.W.H., and R.E.V. wrote the manuscript and all authors critically revised the manuscript.

## Conflicts of Interest Statement

The authors have no conflicts of interest to declare.

## Notes

https://doi.org/10.5281/zenodo.11494629

### Competing Interest Statement

The authors have declared no competing interest.

### Summary of Updates

GEO accession numbers have been removed.

