## Supplementary Information for "Infiltrating lipid-rich macrophage subpopulations identified as a regulator of increasing prostate size in human benign prostatic hyperplasia"

This supplementary information document contains 11 supplementary figures and 3 supplementary tables.

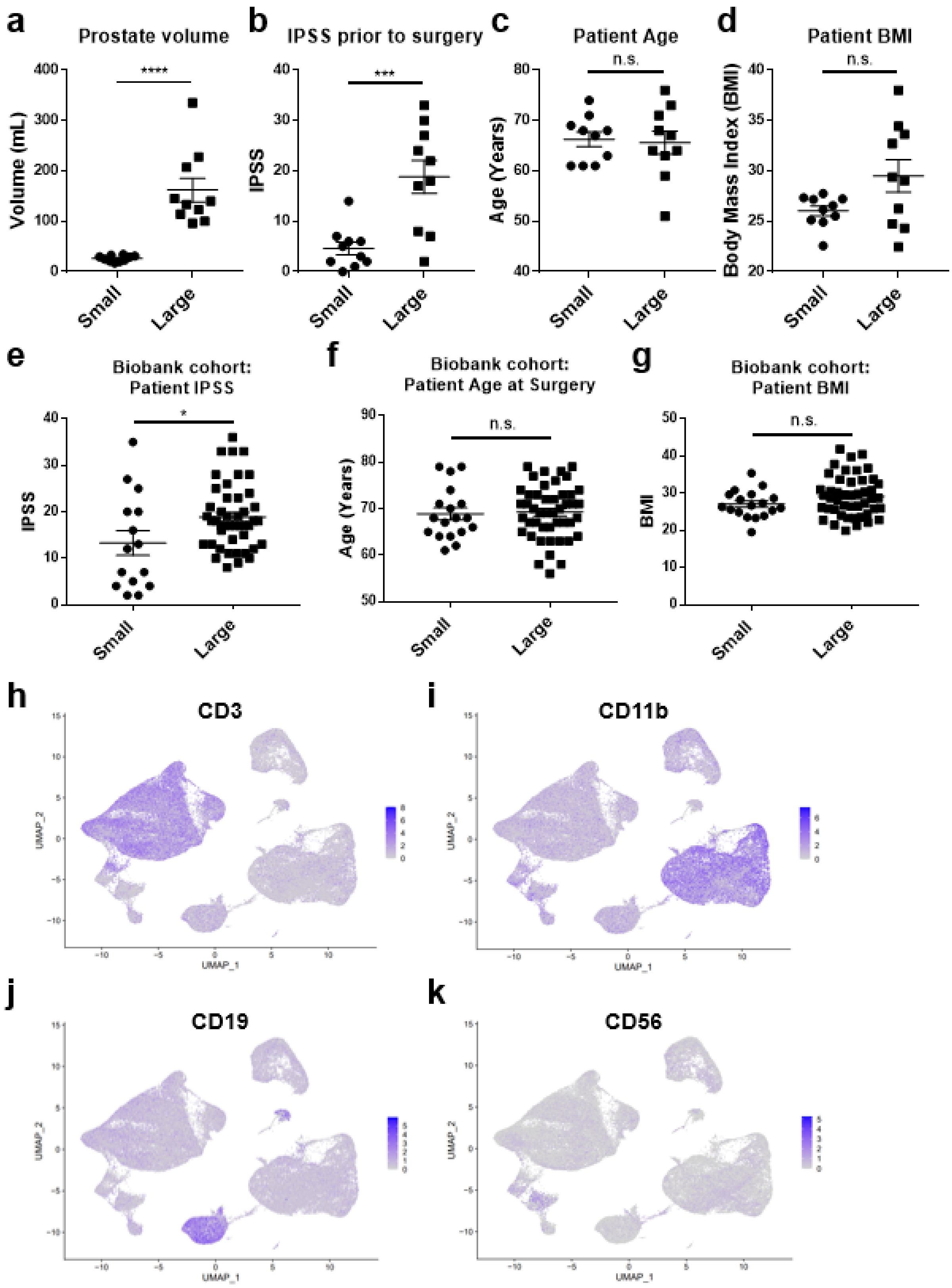
a
Prostate volume
****
Volume (mL)
b
IPSS prior to surgery
***
IPSS
c
Patient Age
n.s.
Age (Years)
d
Patient BMI
n.s.
Body Mass Index (BMI)
Small
Large
e
Biobank cohort:
Patient IPSS
*
IPSS
f
Biobank cohort:
Patient Age at Surgery
n.s.
Age (Years)
g
Biobank cohort:
Patient BMI
n.s.
BMI
h
CD3
i
CD11b
j
CD19
k
CD56
UMAP_1
UMAP_2

**Supplementary Figure S1. scRNA-seq general patient characteristics and identification of major CD45+ cell populations. a)** Prostate volume estimation by TRUS, MRI, or CT scan (****p<0.0001), **b)** patient IPSS prior to surgery (***p=0.0008), **c)** patient age at surgery (p=0.7987), and **d)** patient BMI prior to surgery (p=0.0552) is shown for n=10 patients with small prostates and n=10 patients with large prostates. Statistical analysis was performed using a two-tailed unpaired t-test in (a-d). All patients are biologically independent. Error bars represent the mean +/- SEM for each graph. n.s.=not significant. **e-g)** Analysis of a larger cohort of biobank patients with a clinical diagnosis of BPH. Patients with small predicted prostate size (<40 grams) were compared to patients with large predicted prostate size (>90 grams) for e) Patient IPSS (p=0.0210), f) Patient age at the time of prostate-related surgical procedure (p=0.8249), and g) Patient BMI at the time of surgery (p=0.1388), compared via two-tailed t-test. **h-k)** CITE-seq analysis was conducted on a subset of the 20 patient samples and allowed for the visualization of protein expression in single cells. Feature plots highlight protein expression of h) CD3 (T cells), i) CD11b (myeloid cells), j) CD19 (B cells), and k) CD56 (NK cells), where blue color indicates positive expression and gray color indicates no detectable expression of the indicated protein.

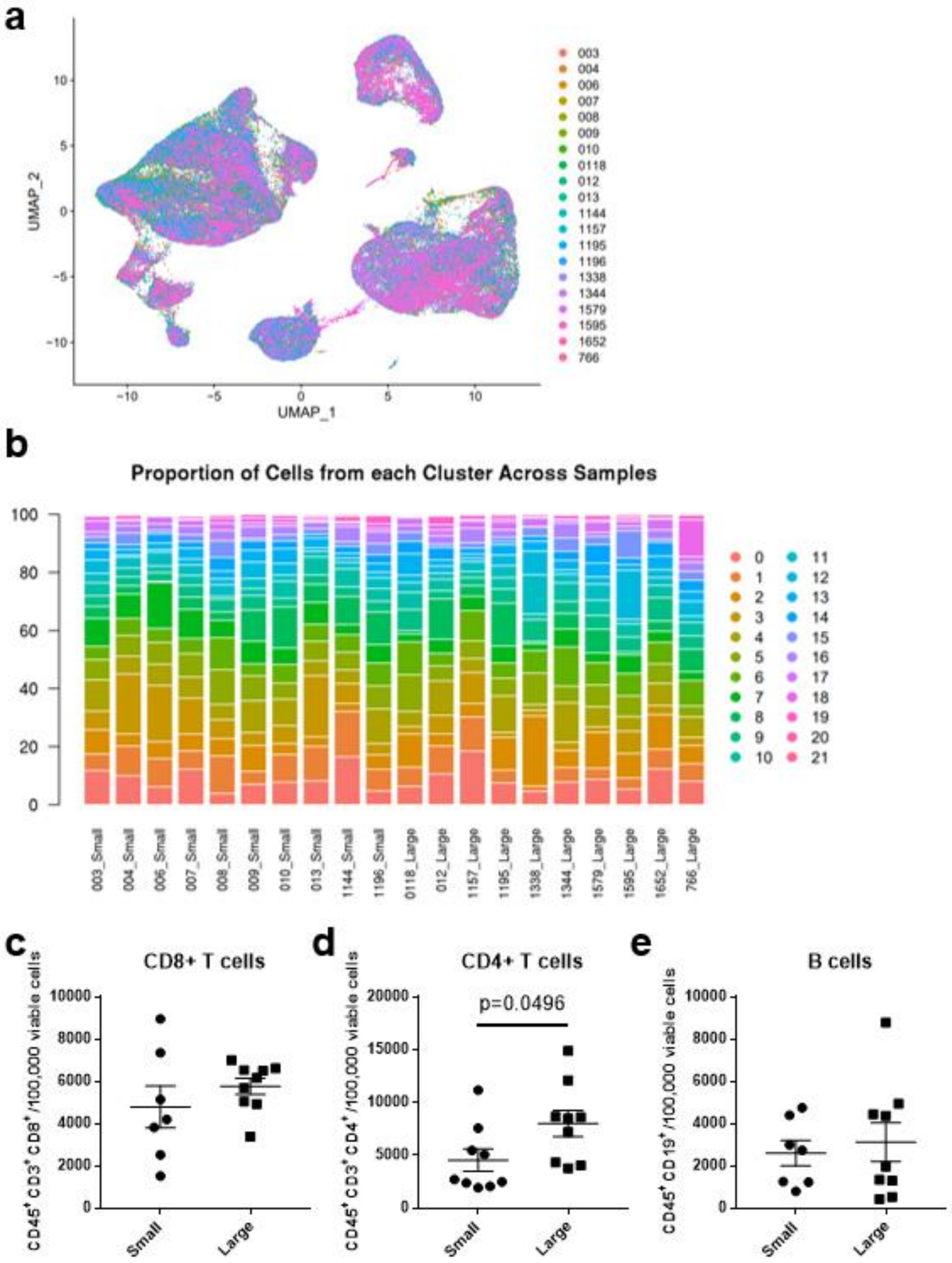


**Supplementary Figure S2. Leukocyte infiltration in human prostate TZ tissue on a per-patient basis.** Analysis of CD45+ leukocyte infiltration was conducted on n=10 small and n=10

large prostate tissues using flow cytometry and scRNA-seq analysis. **a)** UMAP plot of 100,459 BPH-associated immune cells, colored to highlight cells from individual patients. **b)** Graph indicates the proportion of cells in each CD45+ scRNA-seq cluster in individual patients (n=20). Colors indicate each cluster identity. **c-e)** Flow cytometry analysis of digested prostate TZ prior from a subset of the scRNA-seq patient cohort (n=7-9 per group). Graphs show the mean +/- SEM of c) CD45+CD3+CD8+ T cells/100,000 viable cells (p=0.3319), d) CD45+CD3+CD4+ T cells/100,000 viable cells (p=0.0496), and e) CD45+CD19+ B cells/100,000 viable cells (p=0.6593) using a two-tailed unpaired T test. Viable cells were gated zombie violet negative.

a

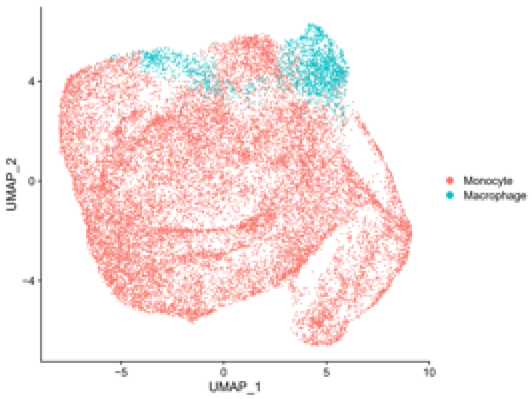
UMAP_2
UMAP_1
Monocyte
Macrophage

b

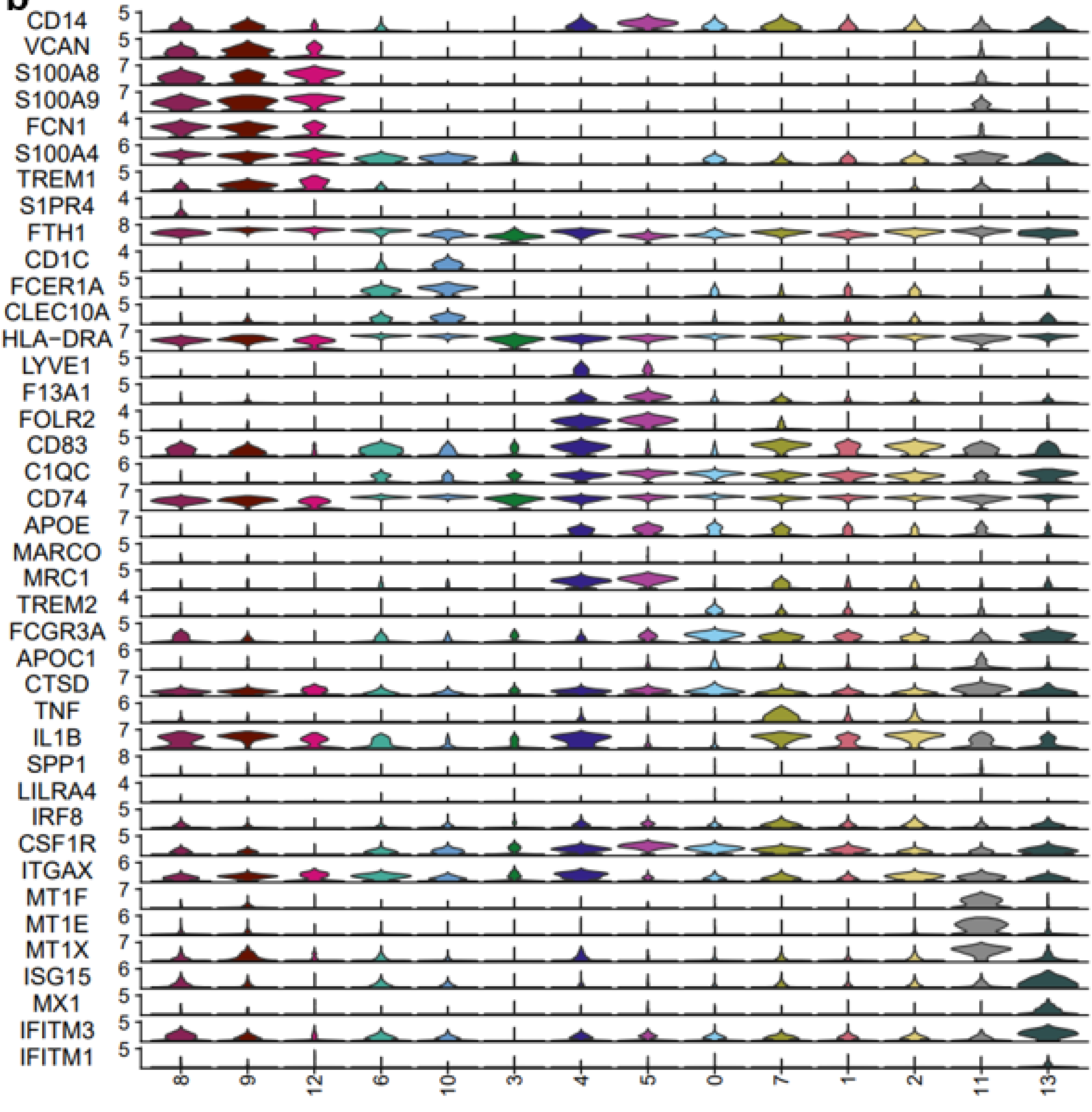
CD14
VCAN
S100A8
S100A9
FCN1
S100A4
TREM1
S1PR4
FTH1
CD1C
FCER1A
CLEC10A
HLA-DRA
LYVE1
F13A1
FOLR2
CD83
C1QC
CD74
APOE
MARCO
MRC1
TREM2
FCGR3A
APOC1
CTSD
TNF
IL1B
SPP1
LILRA4
IRF8
CSF1R
ITGAX
MT1F
MT1E
MT1X
ISG15
MX1
IFITM3
IFITM1
8
9
12
6
10
3
4
5
0
7
1
2
11
13

**Supplementary Figure S3. Identification of myeloid cell subpopulations. Myeloid cells were evaluated via subclustering analysis. a)** UMAP plot indicating the myeloid cell designations using singleR, where individuals cells are labeled as monocytes (pink) or macrophages (blue). **b)** Stacked violin plots of marker genes used to identify myeloid cell subclusters in Figure 2. Clusters are arranged based on proposed similarity.

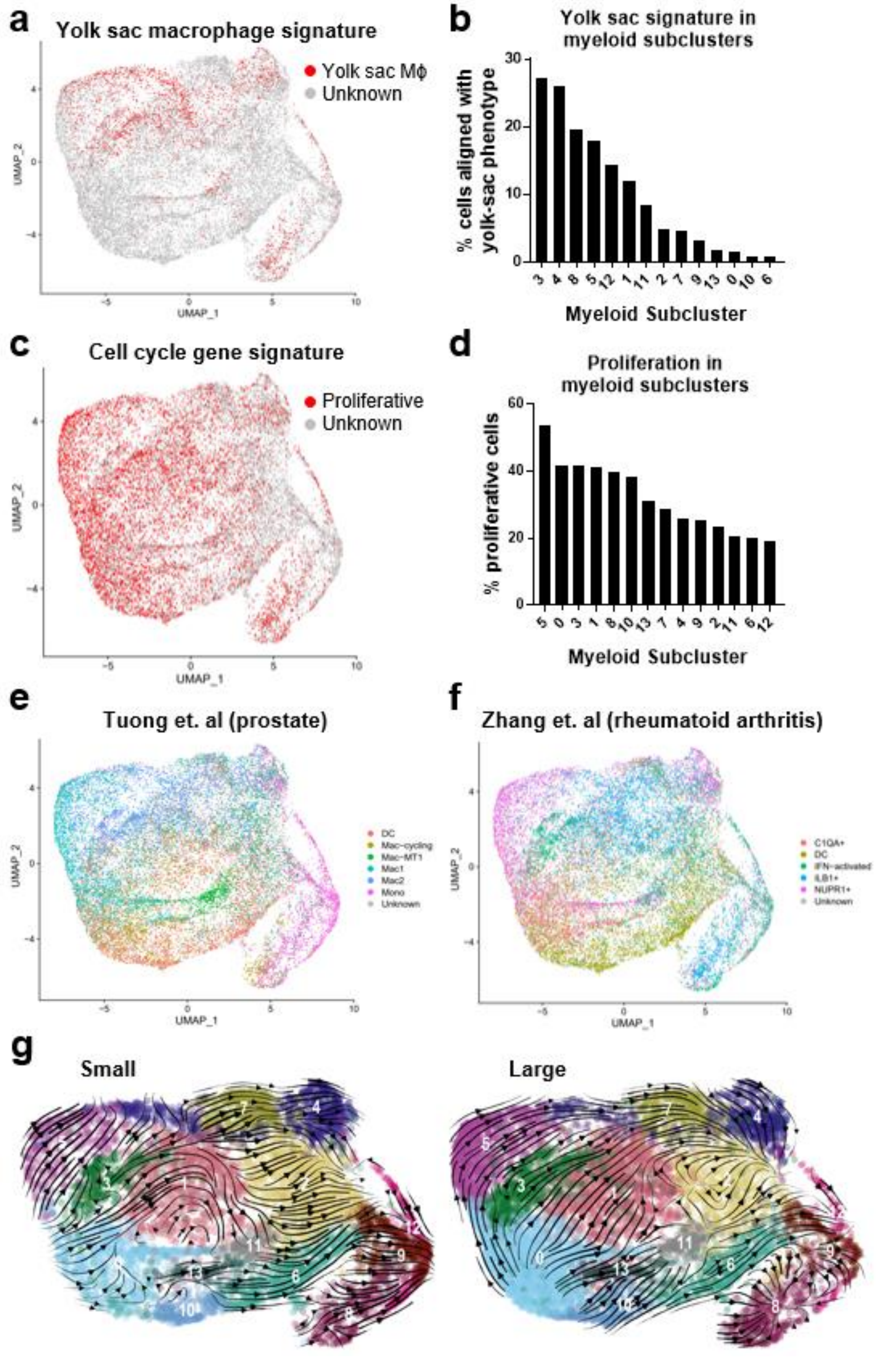
a
Yolk sac macrophage signature
Yolk sac Mϕ
Unknown
UMAP_2
UMAP_1
b
Yolk sac signature in myeloid subclusters
% cells aligned with yolk-sac phenotype
3 4 8 5 12 1 11 2 7 9 13 0 10 6
Myeloid Subcluster
c
Cell cycle gene signature
Proliferative
Unknown
UMAP_2
UMAP_1
d
Proliferation in myeloid subclusters
% proliferative cells
5 0 3 1 8 10 13 7 4 9 2 11 6 12
Myeloid Subcluster
e
Tuong et. al (prostate)
DC
Mac-cycling
Mac-MT1
Mac1
Mac2
Mono
Unknown
UMAP_2
UMAP_1
f
Zhang et. al (rheumatoid arthritis)
C1QA+
DC
IFN-activated
ILB1+
NUPR1+
Unknown
UMAP_2
UMAP_1
g
Small
Large

**Supplementary Figure S4. Additional information relating to myeloid cell subclustering. a-b)** Evaluation of myeloid cell subclustering to determine cells that are transcriptionally similar to yolk sac-derived macrophages was determined via training a classifier using Garnett. **a)** Feature plot indicates cells that express the transcriptional profile of yolk sac macrophages in red. Other cells with unknown identity are labeled gray. **b)** Graph indicates the percentage of cells in each myeloid subcluster that were identified in (a) as being similar to yolk-sac derived macrophages. **c-d)** Evaluation of proliferating BPH myeloid cells was determined via training a classifier using Garnett based on expression of cell cycle genes. **c)** Feature plot indicates expression of cell cycle genes, where a higher red intensity indicates greater expression of cell cycle (proliferation) genes. **d)** Graph indicates the percentage of proliferative cells in each myeloid subcluster based on alignment of cell cycle genes in (c). **e-f)** BPH myeloid cells were assigned to defined to subpopulations in e) prostate tissue and f) synovial tissue from rheumatoid arthritis via Garnett-trained classifiers. The plots indicate the assigned identities projected onto the myeloid subclustering UMAP plot. **g)** RNA velocity analysis was conducted to evaluate cellular trajectories within the myeloid subclustering analysis. The dynamical model of the velocity stream is projected onto the UMAP plot for each small (n=10) and large (n=10) samples.

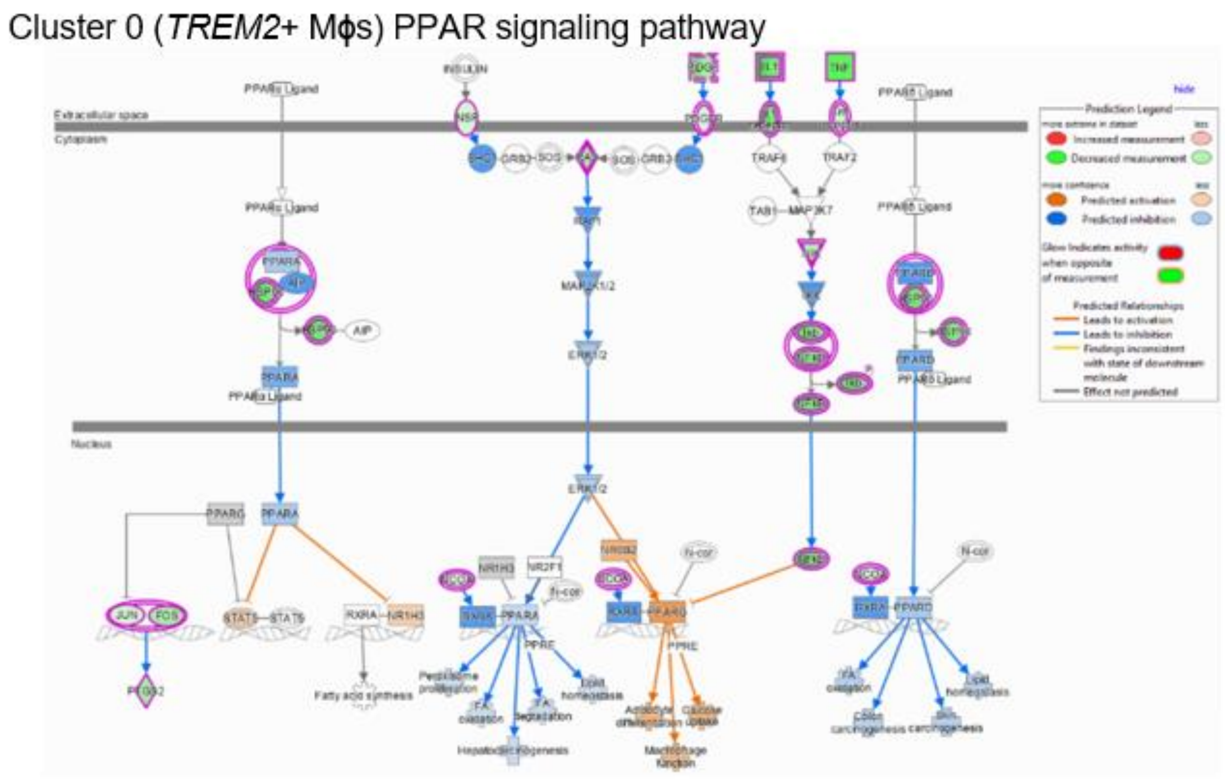


**Supplementary Figure S5. TREM2+ macrophages have activation of the PPAR signaling pathway.** IPA analysis was used to evaluate signaling pathways based on differential gene expression of each subcluster versus all other subclusters. The PPAR signaling pathway for TREM2+macrophage subcluster 0 and predicted activation (orange) or inhibition (blue) is shown.

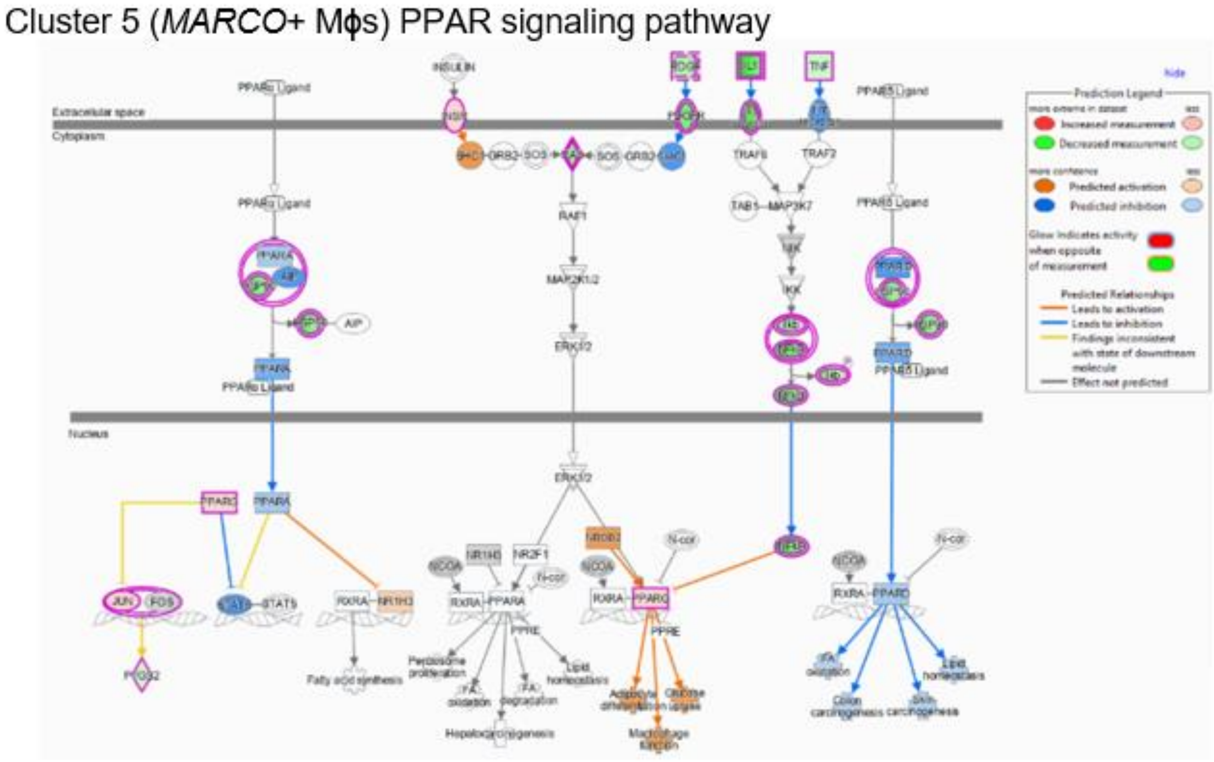


**Supplementary Figure S6. MARCO+ macrophages have activation of the PPAR signaling pathway.** IPA analysis was used to evaluate signaling pathways based on differential gene expression of each subcluster versus all other subclusters. The PPAR signaling pathway for MARCO+ macrophage subcluster 5 and predicted activation (orange) or inhibition (blue) is shown.

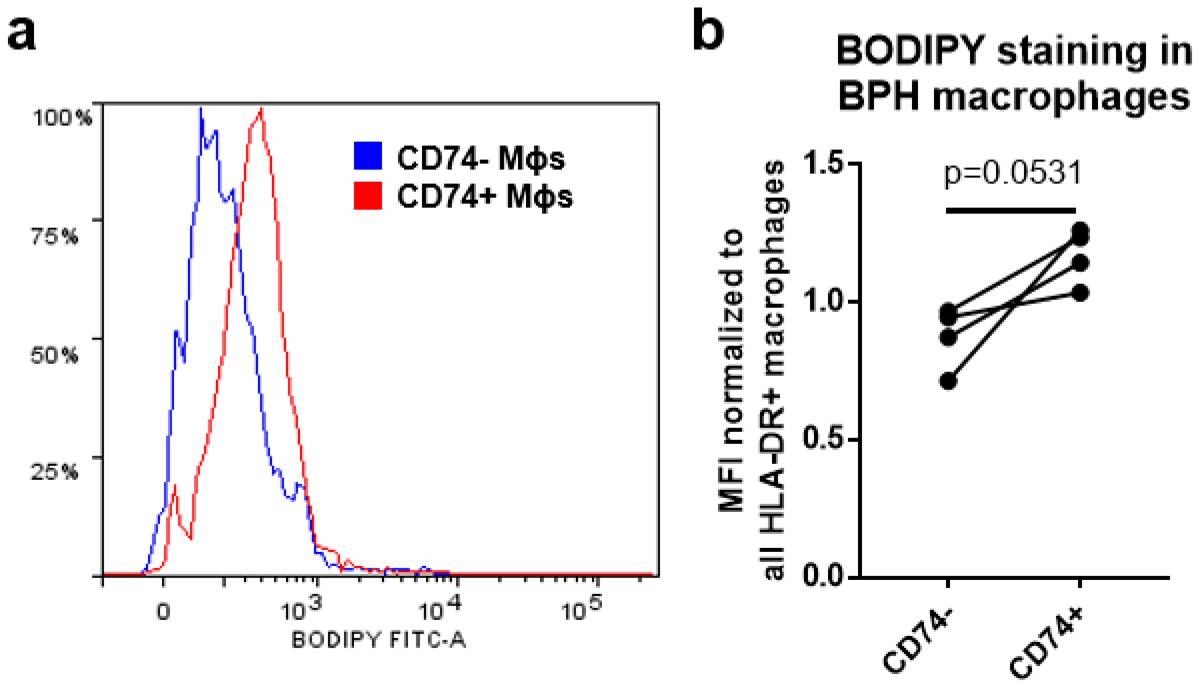


**Supplementary Figure S7. Human prostate transition zone tissues were digested and stained for flow cytometry analysis.** BODIPY staining intensity was measured in CD74+ versus CD74- subpopulations. **a)** Example histogram of BODIPY median fluorescence intensity (MFI) in CD74+ (red) versus CD74- (blue) macrophage subpopulations. **b)** Quantitation of (a) from 4 individual patients, where the BODIPY MFI was normalized to that of all HLA-DR+ macrophages in each sample. p=0.0531 using a paired, two-tailed T test.

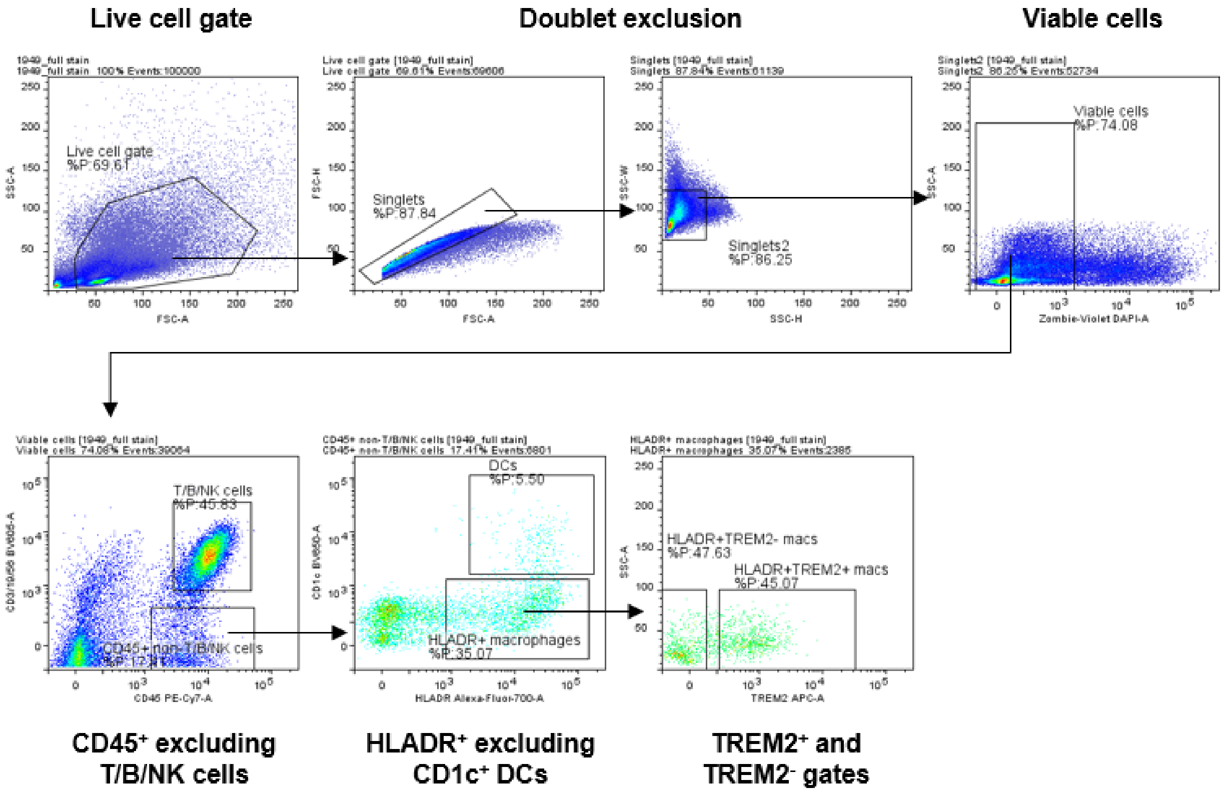


**Supplementary Figure S8. Flow gating strategy for the analysis of TREM2+ versus TREM2- macrophages from human BPH tissues.** The initial live cell gate was drawn on the SSC vs FSC plot, followed by doublet exclusion gates and viable cell gating on zombie violet-negative events. CD45+ cells that were negative for CD3/19/56 were then evaluated for HLA-DR expression. HLADR+CD1c- macrophages were then separated into TREM2+/- populations, where BODIPY staining was individually evaluated and overlaid for analysis.

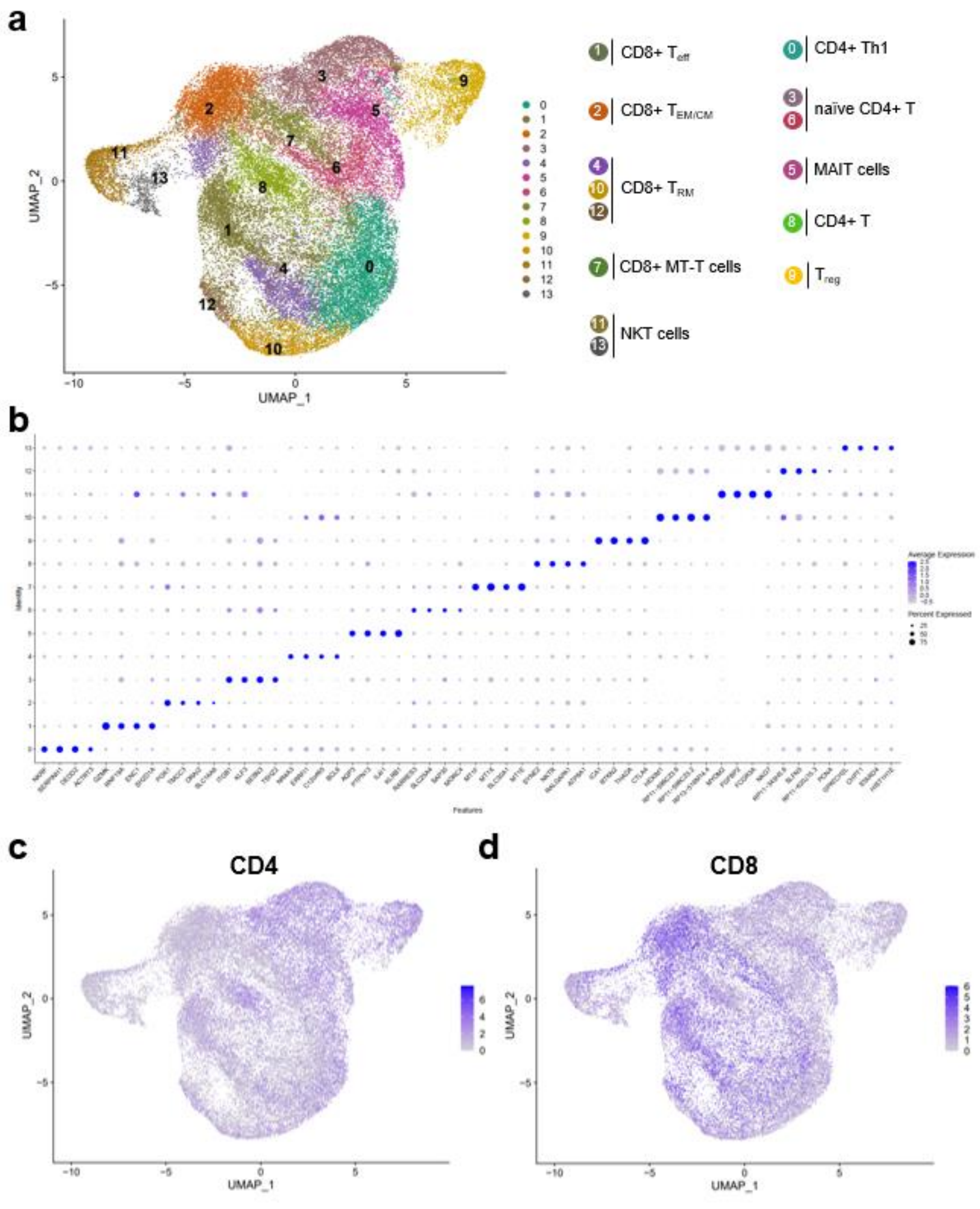


**Supplementary Figure S9. T cell subclustering indicates limited changes among subpopulations in large versus small prostates.** T cells and CD3+ NK cell clusters from scRNA-seq analysis in Figure 1 were used for subclustering analysis after removal of cells expressing keratin genes or myeloid/B cell specific genes. **a)** UMAP plot indicating 14 T/NKT cell subclusters with putative identity. $T_{eff}$=effector T cells, $T_{EM/CM}$=effector memory or central memory T cells, $T_{RM}$=resident memory T cells, MT-T cells = metallothionein-high T cells; MAIT=mucosal-associated invariant T cells, and $T_{reg}$=regulatory T cells. **b)** Dot plot indicating the top 4 marker genes for each cluster, where size of the dot indicates the percentage of cells in the cluster expressing the gene and blue intensity indicates average expression of the gene among all cells in the cluster. **c-d)** CITE-seq analysis was conducted on a subset of the 20 patient samples and allowed for the visualization of protein expression in single cells. Feature plots highlight protein expression of c) CD4 and d) CD8, where blue color indicates positive expression and gray color indicates no detectable expression of the indicated protein. Note that cells with gray color may be from samples that did not include CITE-seq analysis.

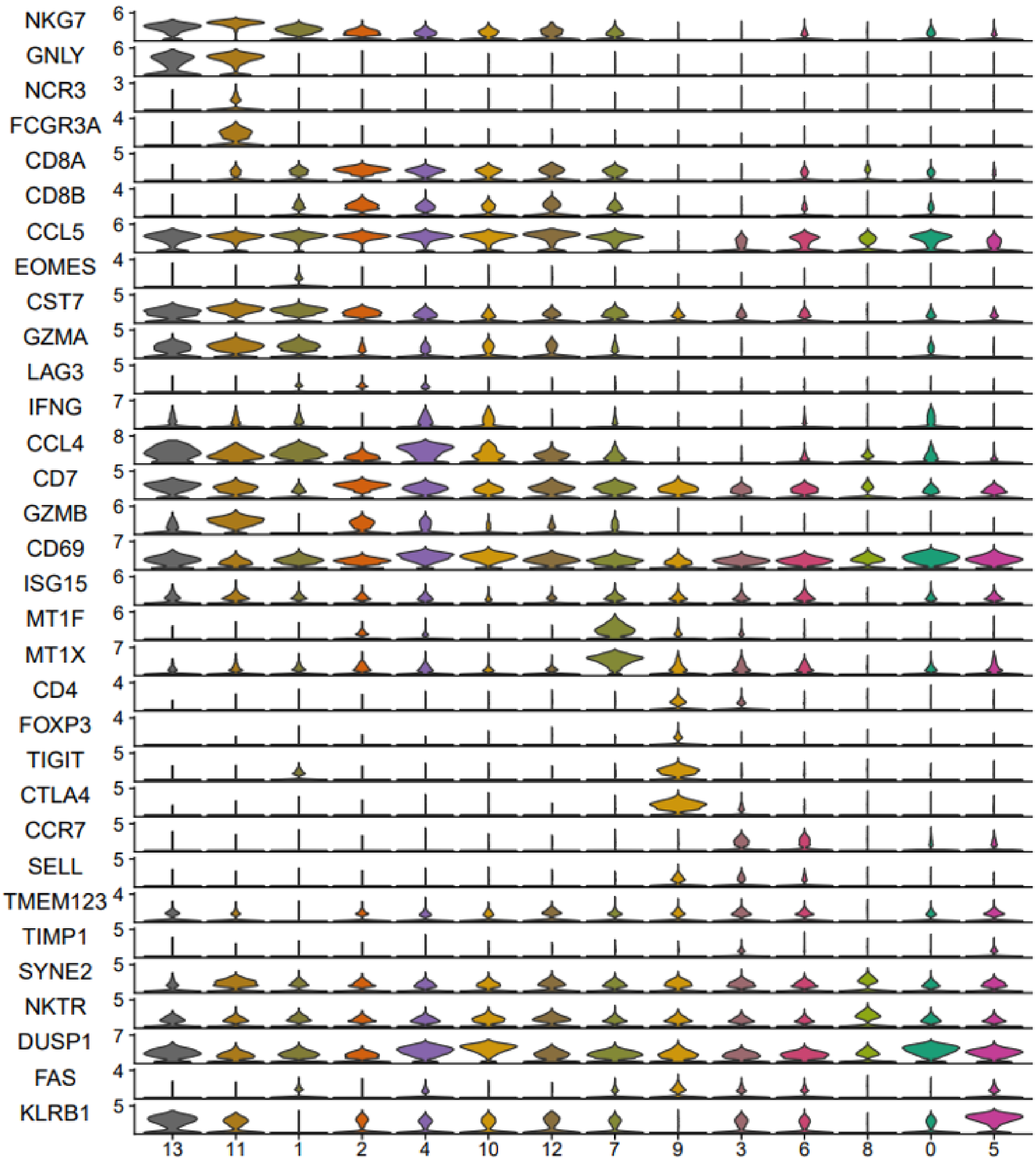


**Supplementary Figure S10. Genes used to identify T cell subpopulations.** Stacked violin plots show gene expression of the various genes used to identify T cell subclusters. Clusters are arranged by proposed similarity.

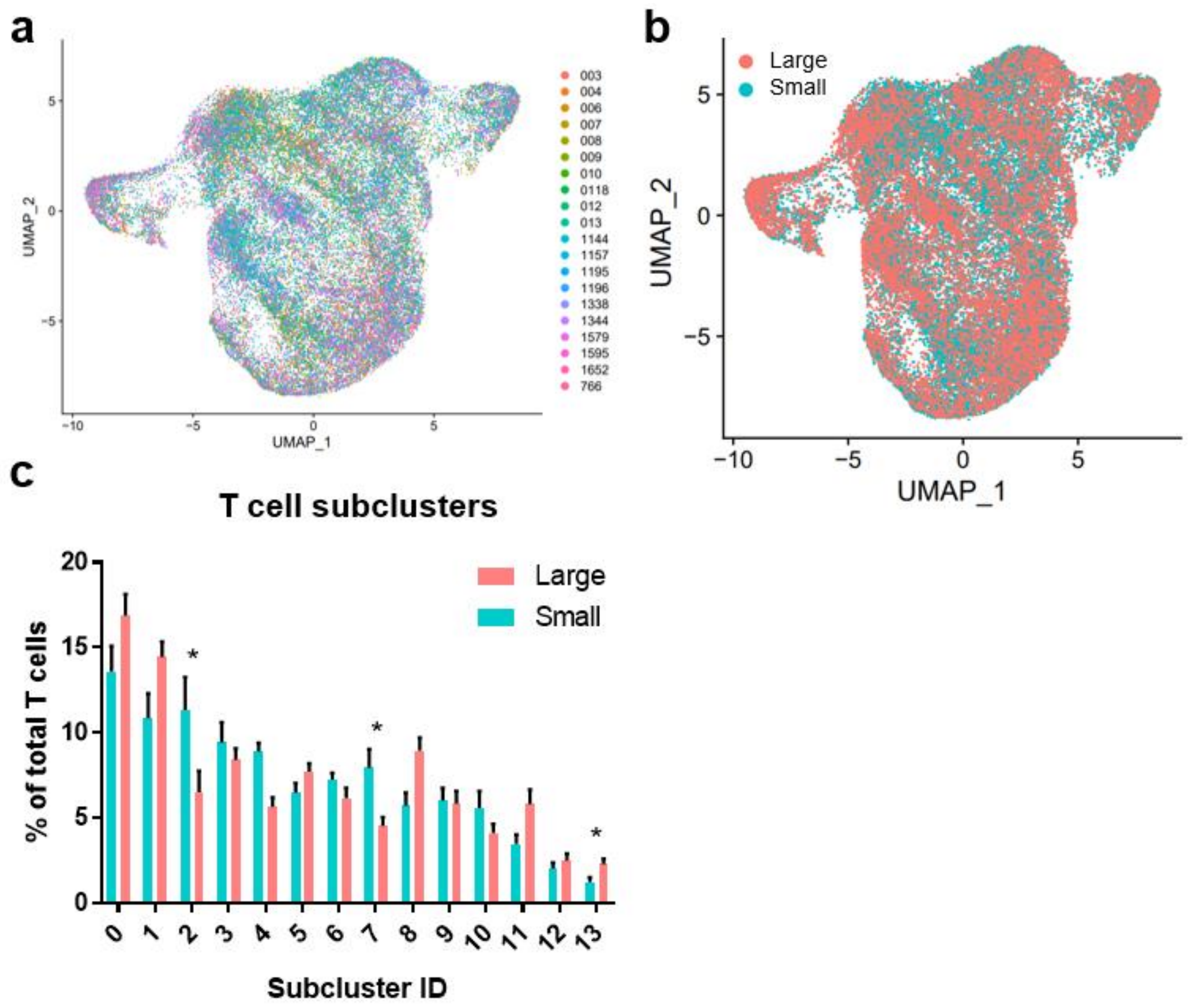


**Supplementary Figure S11. T cell subpopulation changes in large versus small prostate TZ tissues. a)** UMAP plot of T/NKT cell subclustering analysis, colored to highlight cells from individual patients. **b)** UMAP plot of T/NKT subclusters, colored to highlight cells from small (blue) or large (pink) prostates. **c)** Graph representing the mean +/- SEM of the percentage of cells in each T cell subcluster among total T cells between samples in large (pink) versus small (blue) prostates (n=10/group). Asterisks indicate significantly changing populations in large vs small prostate tissues by permutation test (*FDR<0.05).

**Supplementary Table S1. Macrophage polarization signature QUSAGE results.** BPH macrophage subclusters were determined to fit the M1 or M2 signature if the results indicated a positive log(fold-change) and p-value/FDR<0.01.

| Macrophage Subcluster | Pathway Name | log.fold.change | p-value | FDR |
|---|---|---|---|---|
| 0 | M1 | -0.237196696 | 0 | 0 |
| 0 | M2 | -0.14040988 | 0 | 0 |
| 1 | M1 | -0.114730529 | 0 | 0 |
| 1 | M2 | -0.010223622 | 0.124533634 | 0.124533634 |
| 2 | M1 | 0.105220036 | 0 | 0 |
| 2 | M2 | 0.039806782 | 6.23E-08 | 6.23E-08 |
| 3 | M2 | -0.135438606 | 0 | 0 |
| 3 | M1 | -0.070256232 | 0 | 0 |
| 4 | M1 | 0.031018465 | 0.000184514 | 0.000369028 |
| 4 | M2 | 0.016425192 | 0.047234392 | 0.047234392 |
| 5 | M1 | -0.158382662 | 0 | 0 |
| 5 | M2 | 0.017284076 | 0.019256193 | 0.019256193 |
| 6 | M1 | -0.11920269 | 0 | 0 |
| 6 | M2 | 0.141695963 | 0 | 0 |
| 7 | M1 | 0.135516247 | 0 | 0 |
| 7 | M2 | -0.0290059 | 0.003730118 | 0.003730118 |
| 8 | M2 | 0.135956371 | 0 | 0 |
| 8 | M1 | 0.299010676 | 0 | 0 |
| 9 | M1 | 0.429599238 | 0 | 0 |
| 9 | M2 | -0.059009949 | 7.04E-09 | 7.04E-09 |
| 10 | M1 | -0.319631351 | 0 | 0 |
| 10 | M2 | 0.402006665 | 0 | 0 |
| 11 | M2 | -0.078696317 | 0 | 0 |
| 11 | M1 | 0.119197828 | 0 | 0 |
| 12 | M1 | 0.312103101 | 0 | 0 |
| 12 | M2 | -0.038738584 | 0.00106984 | 0.00106984 |
| 13 | M1 | 1.010340694 | 0 | 0 |
| 13 | M2 | -0.018382067 | 0.265059691 | 0.265059691 |

**Supplementary Table S2. Antibodies used for all studies.** Table indicates the antibody target, clone (if monoclonal), manufacturer product number, and dilution used for all antibodies used in these studies.

| Application | Antibody Target | Clone | Species | Isotype | Company | Ref. # | Antibody Dilution |
|---|---|---|---|---|---|---|---|
| FACS | CD45-PE | HI30 | Mouse | IgG1,k | Biolegend | 304058 | 1:20 |
| FACS | EpCAM-APC | 9C4 | Mouse | IgG2b,k | Biolegend | 324208 | 1:20 |
| FACS | CD200-PE/Cy7 | OX-104 | Mouse | IgG1,k | Biolegend | 329212 | 1:20 |
| Flow | CD45-FITC | HI30 | Mouse | IgG1,k | Biolegend | 304006 | 1:20 |
| Flow | CD11b-PE/Cy7 | ICRF44 | Mouse | IgG1,k | Biolegend | 301322 | 1:20 |
| Flow | CD19-APC/Cy7 | HIB19 | Mouse | IgG1,k | Biolegend | 302218 | 1:20 |
| Flow | CD3-APC | UCHT1 | Mouse | IgG1,k | Biolegend | 300412 | 1:20 |
| Flow | CD4-PE | RPA-T4 | Mouse | IgG1,k | Biolegend | 300508 | 1:20 |
| Flow | CD8-BV510 | RPA-T8 | Mouse | IgG1,k | Biolegend | 301048 | 1:20 |
| Flow | TREM2-APC | 237920 | Rat | IgG2B | R&D Systems | FAB17291A | 1:20 |
| Flow | CD3-BV605 | UCHT1 | Mouse | IgG1,k | Biolegend | 300459 | 1:50 |
| Flow | CD19-BV605 | HIB19 | Mouse | IgG1,k | Biolegend | 302243 | 1:50 |
| Flow | CD56-BV605 | 5.1H11 | Mouse | IgG1,k | Biolegend | 362537 | 1:50 |
| Flow | CD45-PE/Cy7 | HI30 | Mouse | IgG1,k | Biolegend | 304051 | 1:50 |
| Flow | HLADR-AF700 | L243 | Mouse | IgG2a,k | Biolegend | 307626 | 1:50 |
| Flow | CD1c-BV650 | L161 | Mouse | IgG1,k | Biolegend | 331541 | 1:50 |
| Flow | CD74-PE | LN2 | Mouse | IgG1,k | Biolegend | 326807 | 1:50 |
| CITE-seq | CD3 | UCHT1 | Mouse | IgG1,k | Biolegend | 300477 | 1:50-1:200 |
| CITE-seq | CD4 | RPA-T4 | Mouse | IgG1,k | Biolegend | 300565 | 1:50-1:200 |
| CITE-seq | CD8 | RPA-T8 | Mouse | IgG1,k | Biolegend | 301069 | 1:50-1:200 |
| CITE-seq | CD11b | ICRF44 | Mouse | IgG1,k | Biolegend | 301357 | 1:50-1:200 |
| CITE-seq | CD19 | HIB19 | Mouse | IgG1,k | Biolegend | 302263 | 1:50-1:200 |
| CITE-seq | CD56 (NCAM) | 5.1H11 | Mouse | IgG1,k | Biolegend | 362561 | 1:50-1:200 |
| IF | TREM2 | 9H4L26 | Rabbit | IgG | ThermoFisher | 702886 | 1:100 |
| IF/IHC | CD68 | KP1 | Mouse | IgG1,k | Dako | M0814 | 1:100 |
| IF | Goat anti-mouse AF594 | Polyclonal | Goat | IgG | Invitrogen | A11032 | 1:2000 |
| IF | Goat anti-rabbit AF488 | Polyclonal | Goat | IgG | ThermoFisher | A27034 | 1:4000 |

**Supplementary Table S3. Run metrics for scRNA-seq of CD45+ cells from human prostate transition zone.** Table includes the run metrics from each scRNA-seq sample, including 10 small and 10 large samples. The metrics indicate high quality data and that the average number of cells captured approached the intended number of 5000 cells.

| **Sample** | **Number of Reads** | **% Q30 Bases in Read** | **% Reads Mapped** | **Estimated Number of Cells** | **Mean Reads per cell** | **Median Genes per cell** |
|---|---|---|---|---|---|---|
| 003_small | 437,303,375 | 91.3 | 96.2 | 6,388 | 68,457 | 1,692 |
| 004_small | 266,767,655 | 91.7 | 96.7 | 4,163 | 64,080 | 1,599 |
| 006_small | 258,138,447 | 91.5 | 96.9 | 6,427 | 40,164 | 1,628 |
| 007_small | 765,500,750 | 90.8 | 96.0 | 4,427 | 172,916 | 1,958 |
| 008_small | 552,744,091 | 93.0 | 98.1 | 5,053 | 109,389 | 1,652 |
| 009_small | 361,879,675 | 93.3 | 96.7 | 3,659 | 98,901 | 1,740 |
| 010_small | 245,628,503 | 91.6 | 96.3 | 3,750 | 65,500 | 1,576 |
| 0118_large | 291,734,653 | 93.5 | 97.1 | 3,803 | 76,712 | 1,801 |
| 012_large | 297,618,482 | 90.2 | 96.7 | 5,826 | 51,084 | 1,526 |
| 013_small | 393,520,430 | 92.6 | 96.7 | 5,230 | 75,242 | 1,476 |
| 1144_small | 394,479,925 | 90.0 | 97.0 | 6,010 | 65,637 | 1,662 |
| 1157_large | 303,893,852 | 90.4 | 96.8 | 6,816 | 44,585 | 1,525 |
| 1195_large | 455,027,712 | 92.4 | 96.6 | 5,377 | 84,624 | 1,591 |
| 1196_small | 224,350,221 | 92.7 | 96.8 | 4,554 | 49,264 | 1,489 |
| 1338_large | 319,810,557 | 93.5 | 96.9 | 6,453 | 49,560 | 1,813 |
| 1344_large | 284,285,224 | 93.6 | 97.0 | 4,245 | 66,969 | 1,475 |
| 1579_large | 363,290,890 | 91.9 | 96.0 | 3,503 | 103,709 | 1,549 |
| 1595_large | 344,446,483 | 91.4 | 96.5 | 6,430 | 53,569 | 779 |
| 1652_large | 323,447,472 | 91.8 | 96.7 | 4,269 | 75,767 | 1,552 |
| 766_large | 368,324,247 | 90.8 | 95.6 | 4,076 | 90,364 | 1,335 |
